# Breakout Rooms, Polling, and Chat, Oh My! The Development and Validation of Online COPUS

**DOI:** 10.1101/2021.07.21.453286

**Authors:** Téa S. Pusey, Andrea Presas Valencia, Adriana Signorini, Petra Kranzfelder

## Abstract

We developed and validated a new classroom observation protocol, Online COPUS (E-COPUS), to measure teaching and learning practices in the online learning environment. We collected COPUS and E-COPUS data from 40 STEM courses before, during the transition, and continuation of emergency remote teaching (ERT). Through weekly discussions among observers, we adjusted six of the original instructor COPUS code descriptions and six of the original student code descriptions to fit the online learning environment. We trained 23 observers to conduct E-COPUS utilizing both in-person and online lecture recordings. To validate E- COPUS, we consulted an expert panel of science educators and education researchers to provide feedback on our code descriptions and complete a matching activity with our E-COPUS code descriptions. We further examined E-COPUS by analyzing the teaching and learning practices of 6 instructors across in-person and online instruction and found that the online functions of breakout rooms, polling, and the chat were utilized to promote active learning activities in the online learning environment. As we prepare for teaching in the future, it is important to have formative assessment tools designed for all course formats to support assessment and improvement of teaching practices in college STEM classrooms.

## Introduction

### Disruptive Events Led to Innovative Pedagogical Approaches

Online teaching is fundamentally distinct from teaching in-person and requires instructors to develop a new set of skills (Davis & Snyder, 2012; Johnston et al., 2005; Juan et al., 2011; Mayer, 2005). For example, effective online instruction requires thoughtful instructional strategies and course design with a variety of assignments and learning activities (Yang, 2017). There are typically two main classroom formats that have been used during online instruction: synchronous and asynchronous (Reinholz et al., 2020; Skylar, 2009). Some instructors may choose to implement a synchronous classroom format (e.g., videoconference call or live online sessions) to encourage student-instructor interactions and group work in the online learning environment (Heiss & Oxley, 2021; Van Heuvelen et al., 2020). However, teaching in a synchronous format does not guarantee student participation; for example, Reinholz et al. (2020) found an overall decrease in student participation in biology classrooms as the class transitioned from in-person to synchronous instruction during emergency remote teaching (ERT). In contrast, instructors may choose an asynchronous format (e.g., recorded lectures, discussion boards, and at-home assignments) if they are concerned about their students’ abilities to attend live online lectures (Van Heuvelen et al., 2020). While asynchronous instruction is an equitable practice for students who cannot attend synchronous lectures, it may reduce some of the student-instructor and student-student interactions that can occur in synchronous lectures (Van Heuvelen et al., 2020). Furthermore, synchronous lectures, which incorporate active learning activities, have been shown to support equity issues, such as less withdrawal rates among underrepresented student groups (Venton & Pompano, 2021b).

As the global pandemic forced instructors to rapidly transition to ERT, instructors adopted new teaching practices to adjust to the circumstances (Rapanta et al., 2020). However, the transition from an in-person to online learning environment mid-course presented many challenges, including maintaining active student engagement (Giordano & Christopher, 2020). As a result, some instructors were not able to implement best teaching practices for online learning (Youmans, 2020), but others approached this challenge with creativity leading to opportunities for classroom pedagogical innovations, including adaptations of active learning strategies.

### Active learning engages students in the learning process, but online learning calls for new assessment tools

Active learning is an evidence-based teaching practice which requires students to engage cognitively and meaningfully with the course materials (Armbruster et al., 2009; Bransford et al., 1999; Chi & Wylie, 2014; Driessen et al., 2020). There are many benefits associated with the implementation of active learning pedagogies (Chickering & Gamson, 1987; Crouch & Mazur, 2001; Freeman et al., 2014; Hake, 1998; Knight & Wood, 2005; Maciejewski, 2016; Ong et al., 2011; Prince, 2004; Ruiz-Primo et al., 2011; Singer & Smith, 2013; Smith et al., 2005; Tomkin et al., 2019) as they are practices that improve learning for all students, particularly persons excluded because of their ethnicity or race, otherwise known as “PEER” (Asai, 2020). Therefore, shifting large numbers of Science, Technology, Engineering, and Mathematics (STEM) faculty to include even small amounts of active learning in their teaching may effectively educate far more students and raise retention of undergraduate STEM students (Owens et al., 2017). Recently, Denaro et al. (2021) noted a national focus on implementing evidence-based teaching practices to improve the quality of STEM education promoted by, among others, the National Research Council (2012), Olson and Riordan (2012), and AAAS (2019).

During the COVID-19 pandemic, there have been a few studies that have documented how active learning has been implemented during online instruction. To recreate active learning activities that were administered before ERT, Tan et al. (2020) utilized the Zoom breakout room function, which creates small videocall rooms within the main meeting, as well as Padlet, a virtual whiteboard. In pre-assigned breakout rooms, students would have discussions facilitated by an instructor or teaching assistant who were present in each breakout room. By asking students to turn their microphone functions on, groups were highly engaged in discussions. Singhal (2020) also utilized breakout rooms when assigning group active learning activities and moved between groups as they worked collaboratively.

Tan et al. (2020) also utilized Poll Everywhere, an online tool for live polling to actively engage students during synchronous instruction. Similarly, Christianson (2020) utilized Socrative, another online tool for live polling, to assign her students group quizzes at the beginning of class. During the administration of the quiz, students used Microsoft Teams to engage in group discussion on the quiz questions. Tan et al. (2020) found that the Zoom chat function, a messaging system within the video call room that allows for participants to send messages to the group or direct messages to each other, was valuable to maintain interactions among faculty, teacher assistants, and undergraduates in the course. Researchers also found that students in the course seemed to respond to more questions and participate more in the chat compared to in-person observations. In a large-enrollment biochemistry course, Dingwall (2020) designed templates for metabolic pathway templates that students could actively fill out during lecture. Students agreed that these templates were useful in the online learning environment because it allowed them to engage in lecture material rather than passively listen to their instructor. Therefore, while there have been studies documenting what tools were successful in implementing active learning activities in the online learning environment (Christianson, 2020; Dingwall, 2020; Singhal, 2020; Tan et al., 2020), studies have not been conclusive about how or what tools to utilize to document the specific teaching and learning practices adapted in the online learning environment.

### Classroom observation protocols

An array of tools have been developed over the past two decades to measure active teaching pedagogies, especially in STEM courses (Eddy et al., 2015; Owens et al., 2017; Sawada et al., 2002; Smith et al., 2013). Self-report surveys alone or in combination with classroom observations is one method to measure the teaching practices of university instructors (AAAS, 2019; Van der Lans et al., 2018). While surveys and interviews are useful in capturing instructor’s personal experiences about implementing active learning, instructors may perceive themselves to be using more active learning pedagogies than they really are in their classrooms (Ebert-May et al., 2011; Van der Lans et al., 2018). In contrast, reliable and validated classroom observation protocols have been developed to objectively support instructors as they implement and reflect on their active learning activities. To name a few, the Practical Observation Rubric To Assess Active Learning (PORTAAL) (Eddy et al., 2015), the Decibel Analysis for Research in Teaching (DART) (Owens et al., 2017), the Reformed Teaching Observation Protocol (RTOP) (Sawada et al., 2002), and the Classroom Observation Protocol for Undergraduate STEM (COPUS) (Smith et al., 2013) provide a way of collecting unbiased data by a trained third-party or an application, like the Generalized Observation and Reflection Platform (GORP) (Tomkin et al., 2019; University of California Davis, 2018; Van der Lans et al., 2018). However, COPUS provides an objective account of the amount of active learning occurring in the classroom (Smith et al., 2013). Also, it has been a useful tool to document in-person active learning activities at different levels, such as at the department (Kranzfelder et al., 2020), the institutional (Smith et al., 2014), and at multi-institutional levels (Akiha et al., 2018; Smith et al., 2013; Stains et al., 2018), as well as to document the impacts of educational initiatives for research (Akiha et al., 2018; Lund et al., 2015; Stains et al., 2018), professional development (Reisner et al., 2020; Tomkin et al., 2019), and tenure, merit, and promotion (Reisner et al., 2020). COPUS findings have also been clustered in different ways to compare results (Denaro et al., 2021) and has been used in combination with other tools, such the Classroom Discourse Observation Protocol (CDOP), (Kranzfelder, Bankers-Fulbright, et al., 2019). Moreover, COPUS results can be offered as an instructor-friendly visual representation documenting the frequency of instructors’ use of active learning practices for different purposes (Kranzfelder et al., 2020; Reisner et al., 2020; Smith et al., 2014).

### Transitioning to E- COPUS during ERT

During ERT, instructors, institutional assessment programs, and biology education researchers faced a problem of not having a reliable, validated classroom observation tool that could be easily implemented online by trained observers to measure active learning. Although adjusting the original COPUS code descriptions to fit the online learning environment may seem like a seamless transition, many universities stopped conducting COPUS during online instruction due to its complexities. For instance, UC Irvine stopped conducting COPUS in the online learning environment because they “were lacking the resources to validate a novel observation protocol in the face of the numerous other COVID-19-related challenges” (personal communication with Brian Sato, 07/20/2021). Additionally, UC San Diego commented: “We had trained undergrads to do the observations and didn’t think we could ask them on the fly to adjust things” (personal communication with Melinda Owens, 07/20/2021). Some institutions utilized COPUS as an assessment tool to support instructors while transitioning to ERT (Clark et al., 2020); however, they did not validate the tool for this new study context (i.e. online learning environment). Therefore, out of necessity, we developed a reliable and validated observation protocol to document online synchronous practices and online functions that instructors may use into their future teaching. By adapting COPUS for the online learning environment, instructors will have the ability to make comparisons between past, present, and future teaching practices as we move to other instructional modalities, such as in-person, Hybrid, or Hybrid-Flexible (HyFlex) (Beatty, 2019). This case study documents the development and validation of Online COPUS (E-COPUS) and showcases an analysis of in-person COPUS data and E-COPUS data.

### Case Study

This study was approved by UC Merced’s Institutional Review Board, and all participating instructors provided informed consent to anonymously participate in the study (Protocol ID UCM2020-3).

### Study context

We conducted this case study in undergraduate and graduate STEM courses at the University of California, Merced (UC Merced), a research-intensive, minority-serving institution (MSI) in the Western United States. This study is part of a larger ongoing research project funded by two research grants, the Howard Hughes Medical Institute Inclusive Excellence (HHMI IE) awarded to the biology program and the National Science Foundation Hispanic Serving Institution (NSF HSI) awarded to the chemistry and biochemistry department, with the goal of understanding, documenting, planning, and enacting meaningful initiatives to improve teaching and student learning at UC Merced.

We collected COPUS data from at least 11 STEM instructors each semester before the transition to ERT (Fall 2018 through Spring 2020), during the transition to ERT (Spring 2020), and/or during the continuation of ERT (Fall 2020 and Spring 2021) (Table 1). To select our instructors for this study, we pulled participants who were involved in a larger institutional study on assessing teaching and discourse practices in college STEM classrooms. To recruit participants for our larger study before the transition to ERT, we identified instructors who: 1) taught either a lower or upper division undergraduate or graduate STEM course, 2) taught a lecture course (excluding laboratory, discussion, or seminar courses), 3) taught the course in-person between two academic years (2018-2019 and 2019-2020) and 4) taught in either the Fall 2018, Spring 2019, Fall 2019, and/or Spring 2020 semesters. To recruit participants during the transition and continuation of ERT, we identified instructors who taught an introductory biology or chemistry course via online synchronous instruction (excluded in-person and asynchronous online instruction). Lund et al. (2015) found that at least two of three successive classroom observations are necessary to adequately characterize an instructor’s teaching practices. Therefore, instructors interested in participating in participation responded to the invitation by providing their course number and three potential observation dates during the semester.

**Table 1.**
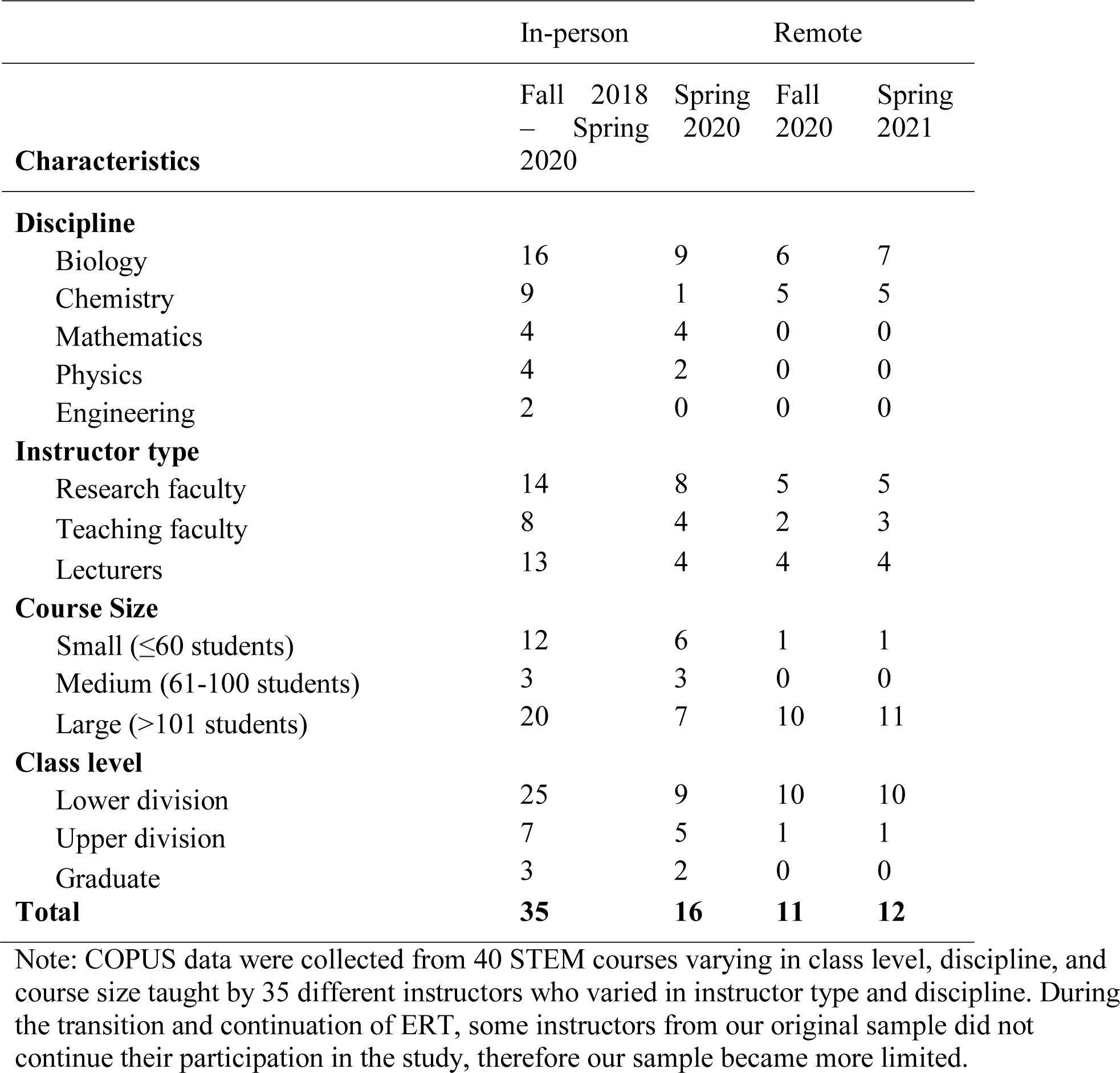
Instructor and course demographics

Instructors taught mainly lower division undergraduate courses from a variety of STEM disciplines, with the majority being in biology or chemistry. All three instructor types from our institution (tenure-track research faculty, tenure-track teaching faculty, and non-tenure track contingent faculty (i.e., lecturers) were observed, with the majority being tenure-track research faculty. Course class sizes ranged from 4 to 292 students, with the mean class size being 110 students. Descriptive information about the instructors and courses included in this study can be found in Table 1.

## Methods

### Data Collection

COPUS documents classroom behaviors in 2-minute intervals throughout the duration of the class session. There are 25 codes in two categories, one to document what the instructor is doing and the other to document what the students are doing. Table 2 offers a description of the COPUS codes: 12 instructors’ behaviors, such as *lecturing* and *moving and guiding*, and 13 student behaviors, such as *listening* and *asking questions*. COPUS provides codes which can be collapsed into many different categories shown in Table 2 (Kranzfelder, Lo, et al., 2019; Smith et al., 2014). Following Smith et al. (2014), we collapsed instructor COPUS codes into four categories: *Presenting, Guiding, Administering,* and *Other*. Following Kranzfelder, Lo, et al. (2019), we collapsed student COPUS codes into four categories: *Receiving, Working & Talking, Assessment, and Other*.

**Table 2.**
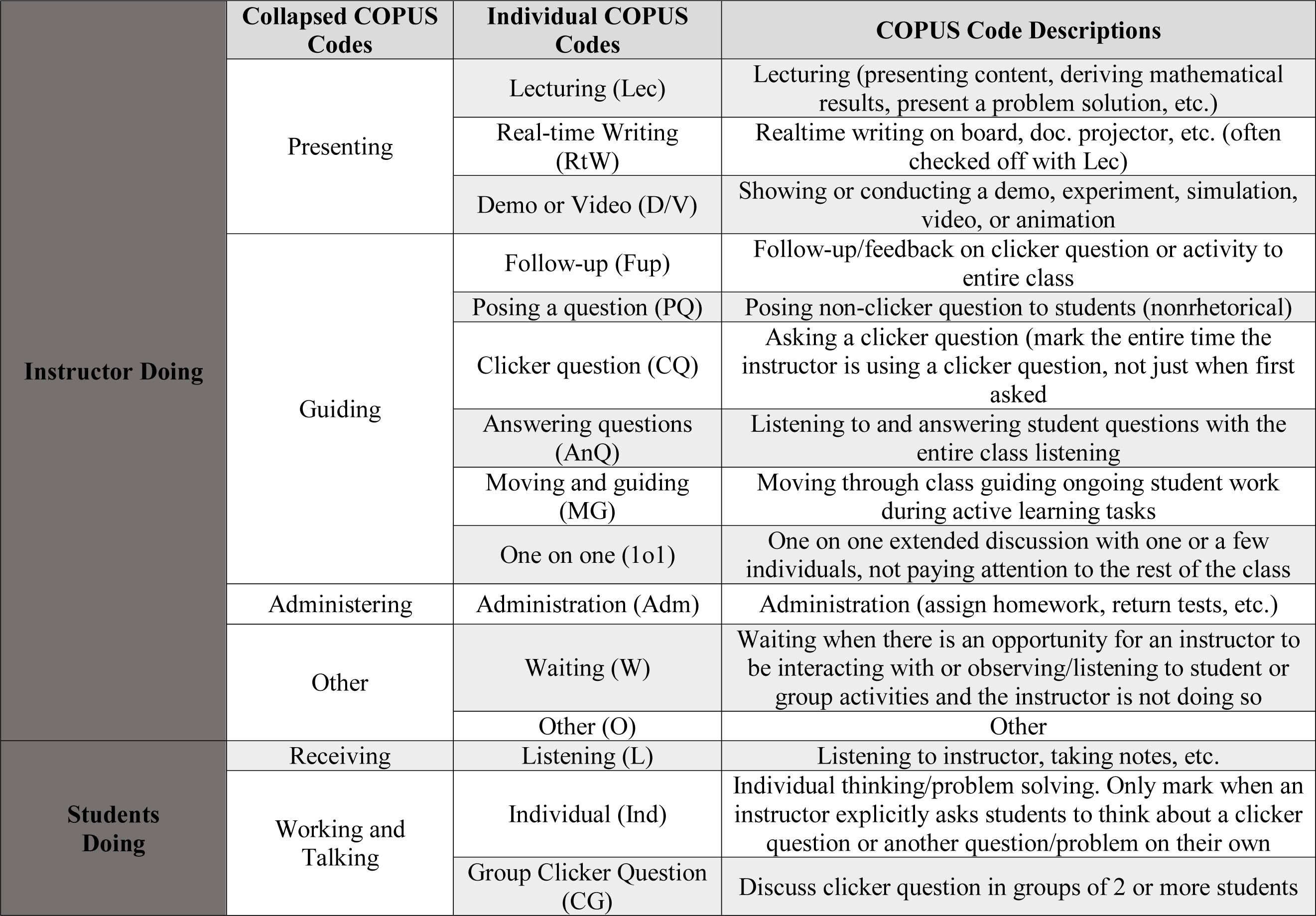
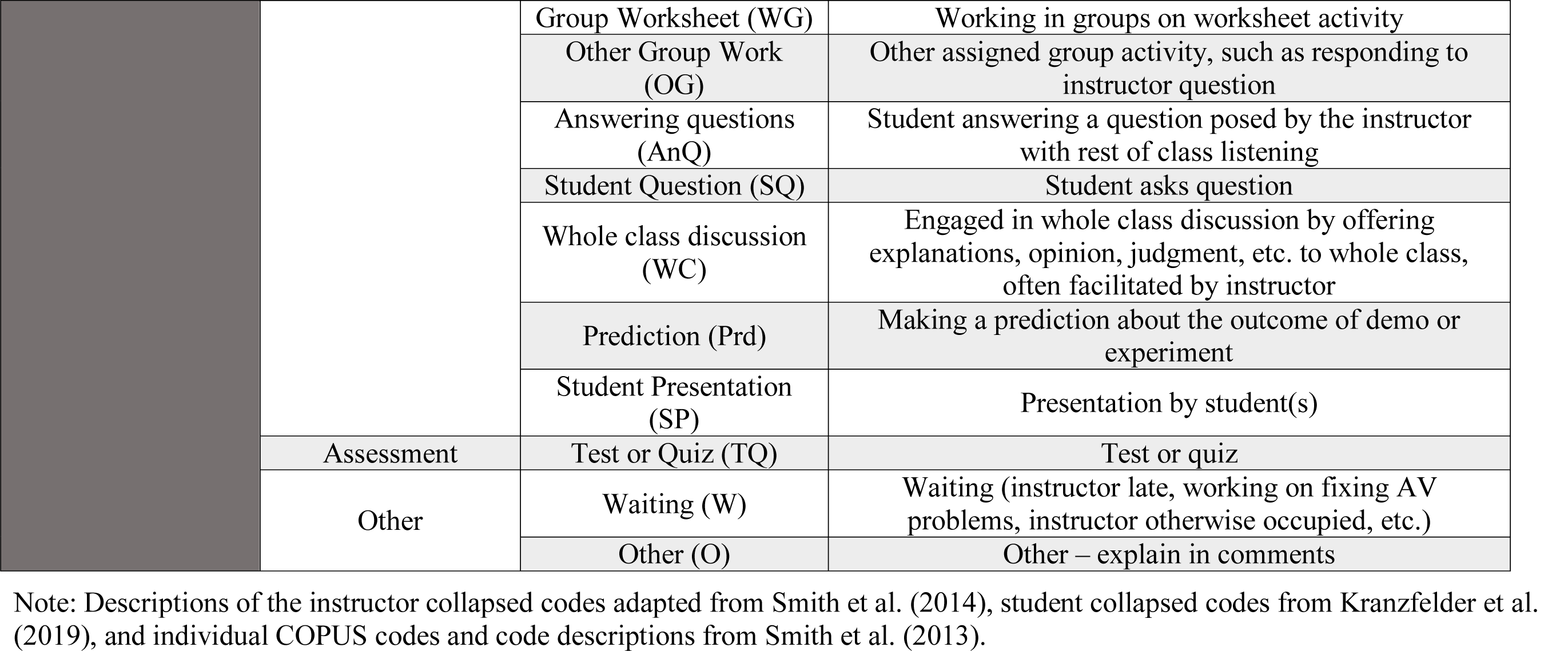
COPUS Coding Scheme.

Once we transitioned to online instruction, observers continued to code classroom observations using in-person COPUS code descriptions, taking detailed notes of instructor behaviors, uses of online functions such as the messaging function, and student behaviors. Following the observation, observers met for up to 30 minutes to discuss codes and resolve any inconsistencies among coders until reaching 100% agreement. During this tabulation, coders also discussed how they coded new online functions or behaviors and their rationale. When coders came to an agreement on how an online function or behavior should be coded, they brought the scenario to our research team’s weekly meeting for group consensus. Once the entire group came to a consensus, the online classroom scenario was added to a shared document among the entire research group to record how we coded that scenario for future reference. After a semester of adapting the original COPUS code descriptions in the online environment, our research team took all the scenarios we had documented and created our finalized online COPUS, or E- COPUS, codebook (File S1).

### Training & Reliability

We trained 15-23 observers for four hours between two training sessions. Each of the two sessions consists of pre- and post-activities as well as a 45-minute coding activity, which utilizes in-person lecture recordings from our home institution (File S2). Additionally, we created a COPUS Frequently Asked Questions (FAQs) to describe in further detail what behaviors we categorize under different COPUS codes and what codes we pair together when a particular behavior occurs (File S3, S4). These training sessions follow an adapted and extended version of the COPUS training in Smith et al. (2013). We adjusted our COPUS training protocol for online instruction (File S2). Our protocol for E-COPUS training contained all the aspects of the in-person COPUS training, except the videos utilized in the coding activity included both online and in-person video recordings from our home institution.

To quantify the degree of agreement between 23 observers using E-COPUS for classroom observations after training, we calculated inter-rater reliability (IRR) using Fleiss’ Kappa at two different points: 1) before coding in-person (File S5) and 2) before coding online (File S6). Landis and Koch (1977) suggest the following interpretations of Fleiss’ Kappa (κ): 0.0-0.20 poor to slight agreement, 0.21-0.40 fair agreement, 0.41-0.60 moderate agreement, 0.61-0.80 substantial agreement, and 0.81-1.0 almost perfect agreement. Before conducting COPUS in-person in Spring 2020, we trained 15 SATAL interns until we reached a moderate kappa average (κ = 0.56,7 95% CI: 0.55-0.56) (File S5). Before conducting E-COPUS in Spring 2021, we trained 23 SATAL interns reached substantial agreement (κ = 0.67, 95% CI: 0.66-0.67) (File S6). In addition to obtaining substantial agreement, SATAL interns met for up to 30 minutes after each E-COPUS observation to discuss codes and resolve any inconsistencies among coders until reaching 100% agreement.

### Validity

To measure content evidence for validity (Yusoff, 2019), we collected expert feedback on E-COPUS by consulting a group of STEM educators and discipline-based education researchers (DBER) at a research-intensive university unrelated to the institution in this study (*n* = 11). The expert feedback panel activities were organized into two parts. First, we (authors TP and APV) collectively presented a subset of the instructor and student codes for the panelists to evaluate the E-COPUS code descriptions. To do this, we showed panelists the original COPUS code descriptions compared to our E-COPUS code descriptions and asked panelists if 1) these behaviors were applicable to them in the online environment, 2) if there were any additional behaviors that they have observed which we should include, and 3) if any of our code descriptions needed further clarification. Additionally, authors AS and PK were present to take notes on the feedback. Our original E-COPUS codebook with the feedback from the expert panel can be found in supplemental materials (File S7). Secondly, we shared online instructor and student codes with the panelists so that they could match them with sample classroom scenarios (File S8). To calculate correct responses for the instructor and student matching activity, we added the number of correct responses for each question and divided the total by the number of questions. For the instructor matching activity, correct responses averaged 85%, while correct responses for the student matching activity averaged 93%. These results provided further proof of content evidence for validity for the online student and instructor codes in typical online classroom scenarios.

Following the expert panel validation, we discussed the feedback provided and made modifications to the E-COPUS code descriptions accordingly. Originally, our E-COPUS codebook contained the original COPUS descriptions (Smith et al., 2013), our research team’s clarifications of the COPUS code descriptions, as well as our E-COPUS code descriptions. First, our expert panel suggested we focus our code descriptions on online behaviors, not clarifications to in-person COPUS codes. We also adjusted the terminology used in the code descriptions to fit multiple meeting applications, such as Skype or Google Meet, so other institutions could adapt our codebook even if they used a different application (Tables 2-3).

**Table 3.** E-COPUS Instructor Coding Scheme

| **Instructor is Doing** | **Individual COPUS Code** | **In-person COPUS Code Description** | **Online COPUS Code Description** |
| --- | --- | --- | --- |
|  | Moving and guiding (MG) | Moving through class guiding ongoing student work during active learning task | Moving through **breakout rooms** guiding ongoing student work during active learning task **or guiding students while they are working on a problem or clicker question (hints/working through a problem) using the microphone or messaging function** |
|  | One-on-one (1o1) | One on one extended discussion with one or a few individuals, giving undivided attention to one or a group of students | One on one extended discussion with one or a few individuals, giving undivided attention to one or a group of students **in a breakout room** |
|  | Posing a question (PQ) | Posing non-clicker question to students (non-rhetorical) and waiting for students to respond | Posing non-clicker question to students (non-rhetorical) **using the microphone or messaging function and** waiting for students to respond |
|  | Answering questions (AnQ) | Listening to and answering student questions with the entire class listening | Listening to and answering student questions **using the microphone or messaging function** with the entire class listening |
|  | Clicker question (CQ) | Asking a clicker question (mark the entire time the instructor is using a clicker question, not just when first asked) | Asking a clicker question **or online poll** (mark the entire time the instructor is using a clicker question, not just when first asked) |
|  | Administration (Adm) | Administration (assign homework, return tests, etc.) | Assigning homework, returning tests, **class announcements/agenda, assign to breakout rooms, etc.), when the instructor is waiting for students to answer a non-clicker question (i.e., think-pair-share), or administering a test or quiz** |

Note: Descriptions of the in-person COPUS code descriptions adapted from Smith et al. (2013). Modifications to online COPUS code descriptions are noted in bold.

### Data Analyses

To understand the differences in instructional practices and student behaviors in the online environment, we analyzed six instructors who were observed during in-person (Fall 2019) and continuation of ERT (Fall 2020) (Figures 1-2). We developed pseudonyms that represented the race and gender for each instructor to keep the instructors de-identified, but still reflect their demographics. We used the collapsed categories developed by Smith et al. (2014) to compare instructor’s in-person COPUS to the E-COPUS data and Kranzfelder, Lo, et al. (2019) to compare student’s in-person COPUS to the E-COPUS data. To determine any differences among the collapsed COPUS codes by class format, we followed Smith et al. (2014), Lewin et al. (2016), and Kranzfelder et al. (2020) to examine the average percentage of COPUS codes implemented between each instructor’s in-person and online class sessions. To do this, we took the sum of a singular COPUS/E-COPUS code (i.e., *lecturing)* and divided it by the total number of codes that appeared within the class session.

**Figure 1.**
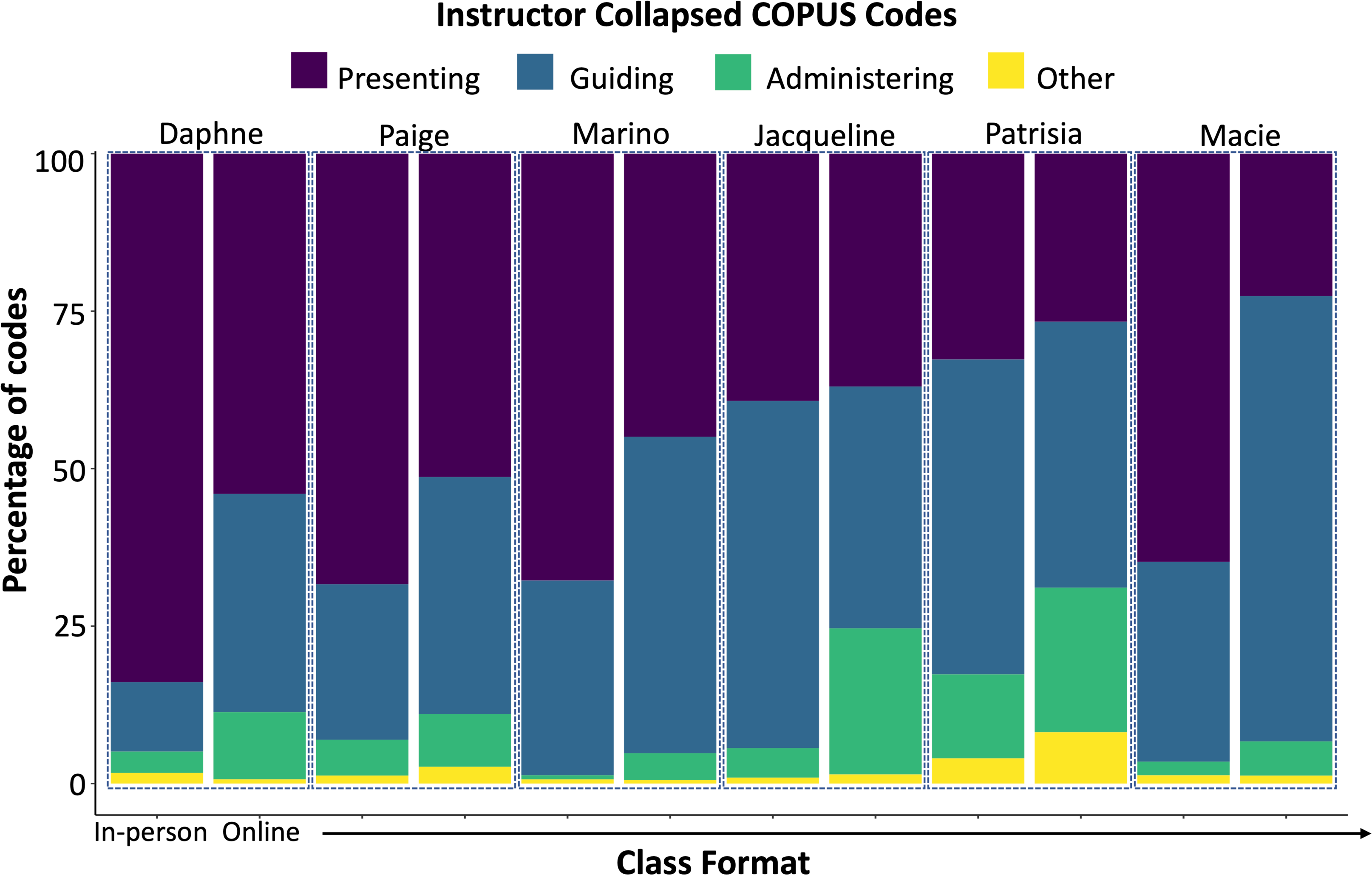
A comparison of the average percentage of collapsed instructor COPUS codes of three different class sessions from six STEM instructors during in-person and online instruction.

**Figure 2.**
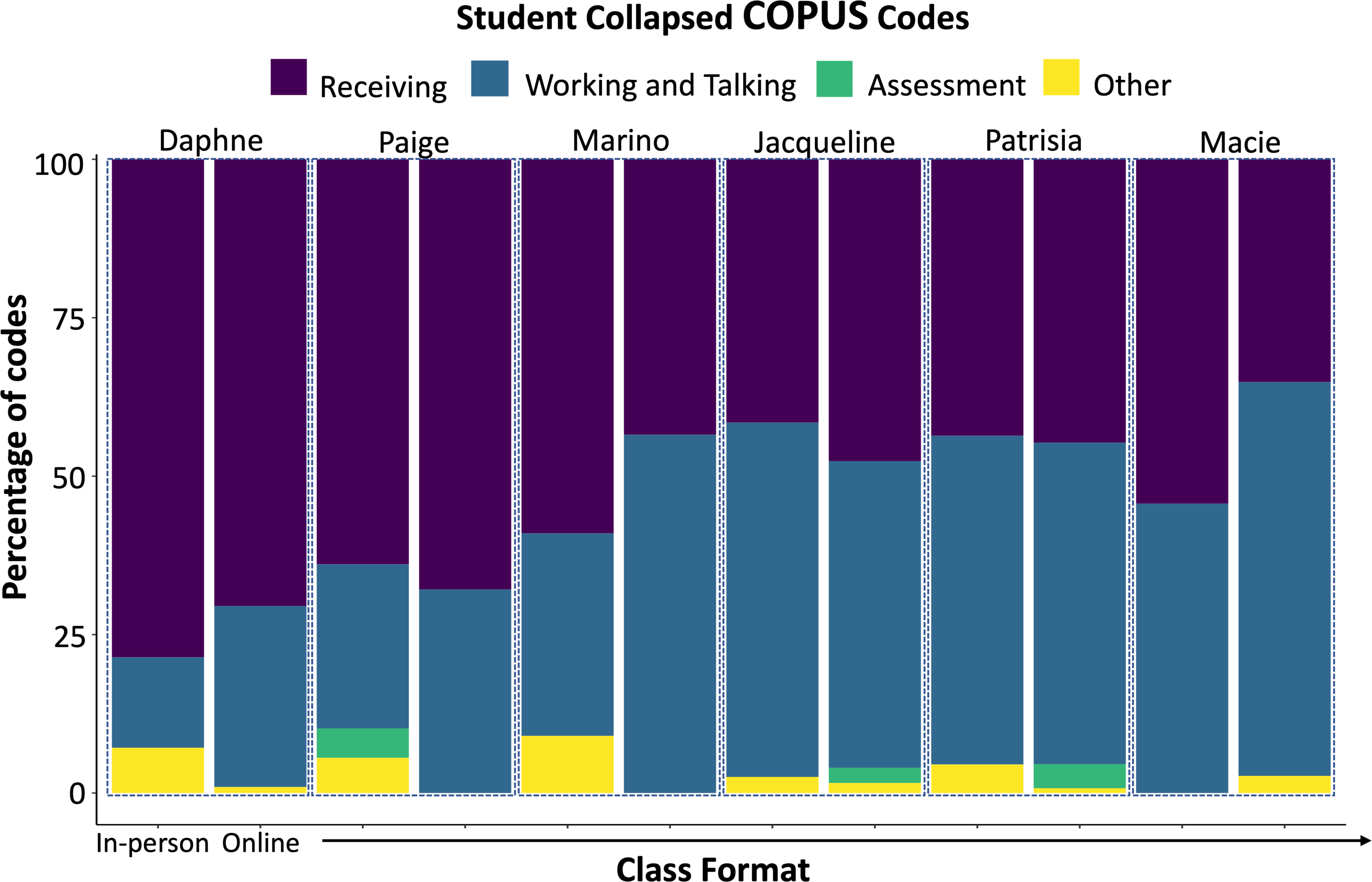
A comparison of the average percentage of collapsed student COPUS codes of three different class sessions from six STEM instructors during in-person and online instruction.

To understand how the online learning environment impacted specific instructor and student behaviors, we compared Macie’s individual in-person and online instructor and student behaviors (Figure 3) using the finalized E-COPUS coding scheme (Tables 2-3). Macie was observed in Fall 2019 (in-person) and in Spring 2021 (online). We selected Macie’s class sessions for analysis because they had the highest increase in instructor *Guiding* between in-person and online instruction as shown in Figure 1. To compare Macie’s individual COPUS and E-COPUS codes, we took the number of two-minute time intervals marked for each individual code and divided it by the total of two-minute time intervals for the class session (Kranzfelder et al., 2020; Lewin et al., 2016; Lund et al., 2015). This calculation overestimates the time spent on a particular code as the behavior is counted for the entire two-minute interval even if the instructor only spends a few seconds on it.

**Figure 3.**
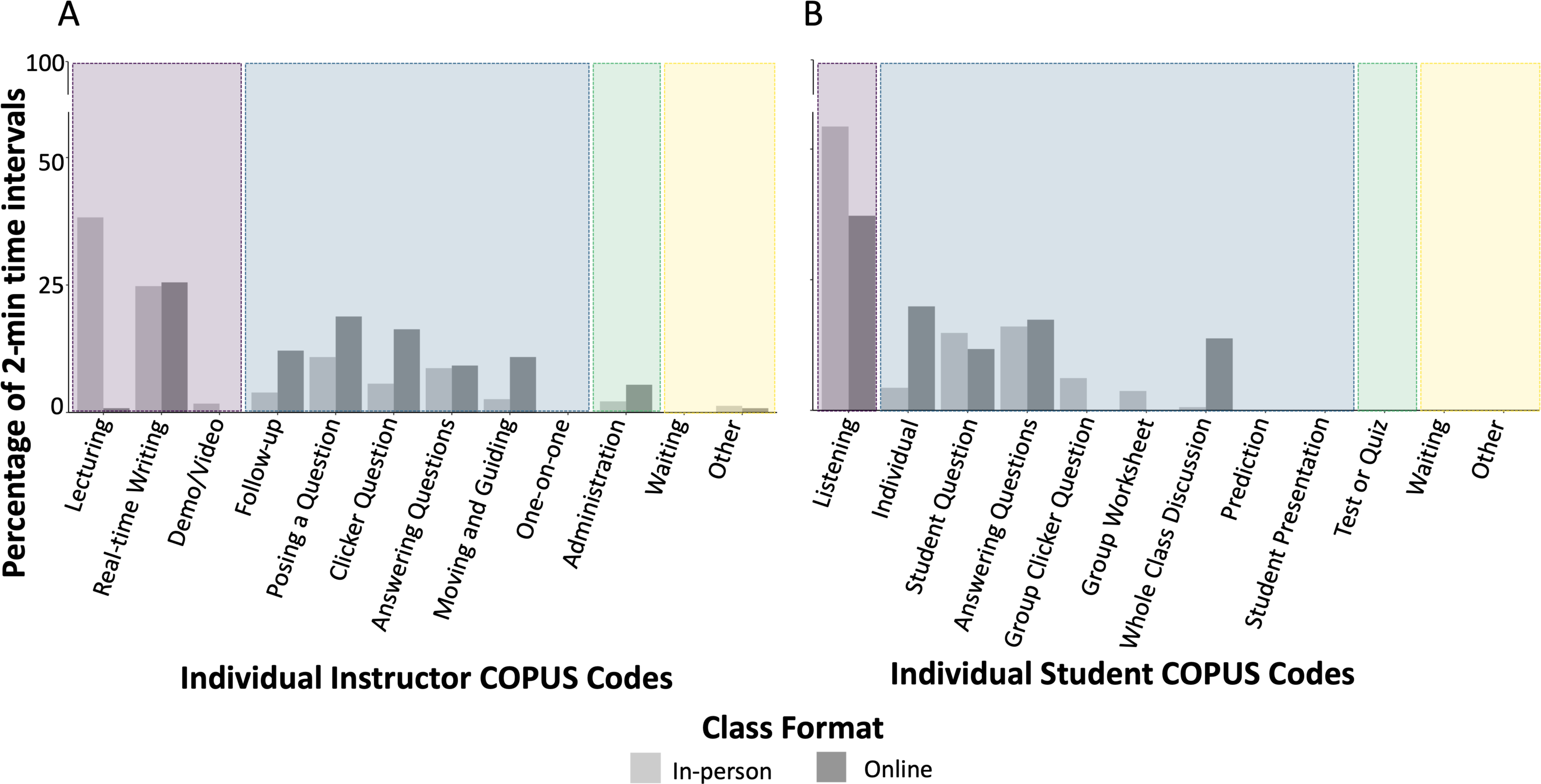
A comparison of the average percentage of two-minute time intervals spent on individual instructor and student COPUS/E-COPUS codes from one STEM instructor (Macie) across three different class sessions during in-person and online instruction. The collapsed student and instructor COPUS codes are color coded following Figures 1-2 for comparison.

## Results

### Instructor E-COPUS Code Descriptions

We adjusted six of the twelve original instructor COPUS code descriptions to document the teaching behaviors we observed in the online learning environment as illustrated in Table 3. We did not change any of the original COPUS codes, but rather adjusted their descriptions to fit the online learning environment. Most adjustments to the in-person COPUS code descriptions were the result of new online functions, such as breakout rooms, polling, and the chat. Tables 3 and 4 present E-COPUS codes and code descriptions for instructor and student behaviors, respectively. To make our E-COPUS code descriptions more inclusive for other meeting software programs, we used the terms “messaging function” and “chat function” interchangeably. See Supplementary Materials for full descriptions of the codebook, which includes inclusion and exclusion criteria for instructor and student behaviors (File S9-S10).

### Breakout Rooms

#### Moving and Guiding (MG)

We changed the code description for *moving and guiding* by adding newly observed behaviors as well as excluded behaviors that no longer fit in the online learning environment. In the online environment, instructors could no longer physically move around the classroom, so we utilized this code when the instructor moved in and out of breakout rooms and guided students in their active learning activity. We also found instructors engaged in *moving and guiding* behaviors without having to move throughout breakout rooms. For example, we also coded *moving and guiding* when an instructor assigned an active learning activity and provided students hints to answer a problem or showed students how to solve the problem as they were working on it. We agreed that this was also a *moving and guiding* behavior even though the instructor did not create breakout rooms because they were still guiding students in an active learning activity (Table 3).

#### One-on-One (1o1)

We changed the code description for *one-on-one* in the online learning environment. Specifically, it occurred when the instructor was moving between breakout rooms and staying with one group for an extended period of time. This behavior would be similar to the instructor walking around the classroom and spending extended time with student groups during group work.

#### Administration (Adm)

We adjusted the description of *administration* to include scenarios that we frequently encountered in the online learning environment, like assigning breakout rooms or assigning an individual thinking question that was not a *clicker question*, such as a think-pair-share. While these behaviors could be interpreted in the “etc.” of the original description, we included them to ensure consensus of coding these behaviors during observations.

### Polling

#### Clicker Question (CQ)

Next, for the *clicker question* code description, we added online functions that appeared in the online learning environment. The most prominent online activity we observed were online polls, such as those used on Zoom or third-party sites, such as Poll Everywhere, Socrative, or Mentimeter, which we coded as a *clicker question*. While not identical, online polls allowed students to individually think and submit their answer to a multiple-choice question as well as see the distribution of student responses, like a *clicker question*.

### Chat

#### Posing a Question (PQ) and Instructor Answering Questions (AnQ)

Lastly, for the codes *posing a question* and *answering questions,* we found that the chat function allowed instructors to ask and answer questions in two ways: verbally or written through the chat. Therefore, we slightly changed these code descriptions to include both modalities.

#### Student E-COPUS Code Descriptions

We adjusted six of the thirteen COPUS code descriptions to document the teaching behaviors we observed in the online learning environment as illustrated in Table 4. While most student codes were easily adaptable to the online environment, some codes required more adjustments. Most of these code descriptions were adapted to include the online functions used in online instruction, such as the chat, as well as any new behaviors that emerged because of the implementation of these functions.

**Table 4.** E-COPUS Student Coding Scheme

| **Student is Doing** | **Individual COPUS Code** | **In-person COPUS Code Description** | **Online COPUS Code Description** |
| --- | --- | --- | --- |
|  | Answering questions (AnQ) | Student answering a question posed by the instructor with rest of class listening | Student answering a question posed by the instructor **using the microphone function, reaction function, annotating function, or messaging function and the instructor acknowledges the answer with the rest of the class listening or student answering other students' questions using the messaging function** |
|  | Whole class discussion (WC) | Engaged in whole class discussion by offering explanations, opinion, judgment, etc. to whole class, often facilitated by instructor | Instructor poses a question or facilitate a whole class discussion in which 2 or more students answer **verbally, using messaging function, or drawing function while the rest of the class is listening** |
|  | Group Clicker Question (CG) | Discuss clicker question in groups of 2 or more students | Discussing clicker question in groups of 2 or more students **in breakout rooms or messaging function** |
|  | Group Worksheet (WG) | Working in groups on worksheet activity | Working in groups of 2 or more students on worksheet activity **in breakout rooms or messaging function** |
|  | Other Group Work (OG) | Other assigned group activity, such as responding to instructor question | Working in groups of 2 or more students on other assigned group activity, such as responding to instructor question **in breakout rooms in breakout rooms or messaging function** |
|  | Student Question (SQ) | Student asks question | Student asks question **using the microphone or messaging function** |

Note: Descriptions of the in-person COPUS code descriptions adapted from Smith et al. (2013). Modifications to online COPUS code descriptions are noted in bold.

### Breakout Rooms

#### Group Clicker Question, Group Worksheet, and Other Group Work (CG, WG, and OG)

In the online learning environment, we found group work could be seen in two ways: 1) when the instructor assigned students to work on an active learning activity in breakout rooms, or 2) when students engaged in group work by discussing an active learning activity in the chat without instructor facilitation. For example, in one observation a group of five students used the chat to work on a clicker question together without any instructor intervention. Since this discussion was not facilitated by the instructor, we concluded it was not a *whole class discussion*, but instead a *group clicker question*.

### Chat

#### Student Answering Questions (AnQ)

The code description for *answering questions* includes all the ways that students could answer a question in the online learning environment. The first and most direct way a student could answer a question posed by the instructor was by responding verbally while the rest of the class was listening. The second way a student could answer a question posed by the instructor was by using the chat function, available for everyone in the class to read, which is like the rest of the class listening as the original code stated. However, in some observations, we noticed that some students’ responses in the chat were unnoticed by the instructor. Furthermore, while it was possible for students to answer an instructor’s question through private messaging, observers were unable to see these responses. Therefore, the description to the code *answering questions* was adjusted to explicitly state “student answering a question posed by the instructor using the microphone or chat function *and* the instructor acknowledges the answer with the rest of the class listening.” Additionally, we noticed that throughout the class session some students would ask and respond to each other’s questions, sometimes without the instructor’s intervention. To acknowledge that these students received an answer to their questions, we deemed it appropriate to code *answering questions* and added “or student answering another students’ question using the chat function” to the code description.

#### Whole Class Discussion (WC)

The online learning environment allowed for students to be involved in a *whole class discussion* utilizing different functions, including the chat, writing, or drawing function. If multiple students answered an instructor’s question using the chat, writing, or drawing function, then we coded *whole class discussion.* For example, if the instructor asked the class to use the drawing function to draw a cell structure on a slide, then observers coded it as *whole class discussion*. Another example of a *whole class discussion* would be if the instructor posed a question the class and multiple students responded in the chat.

#### Student Question (SQ)

The description for the code *student question* was slightly altered to account for the modalities which a student could ask a question in the online environment: verbally or through the chat function. Like *answering questions,* students could ask the instructor questions through the private chat; however, observers are unable to see the behaviors unless the instructor explicitly acknowledges they received a private message with a question. Additionally, the original code description for *student question* did not specify that the whole class must be listening, unlike the original code description for *answering questions*. Therefore, regardless of whether the instructor acknowledged a students’ question, we used *student question* to code this behavior.

### All instructors implemented less *Presenting* practices and more *Administering* during online instruction

We found that all six instructors on average *Presented* less and *Administered* more in the online learning environment (Figure 1). For example, during in-person instruction, most of Macie’s class consisted of *Presenting* (65%), while including some *Guiding* practices (32%) and very little time spent *Administering* (2.2%). However, during online instruction, Macie’s teaching behaviors almost inverted; most of her behaviors were now *Guiding* practices (71%) with some *Presenting* (23%) and a bit more *Administering* (5.5%) practices. Additionally, most instructors (67%) had a higher average of *Guiding* practices during online instruction. However, this was not true for all our instructors. During in-person instruction, most of Jaqueline’s behaviors were *Guiding* (55%) and some *Presenting* (39%). In the online learning environment, Jaqueline used almost as much *Presenting* (37%) practices as she did *Guiding* (38%) practices. This tells us that our instructors adapted to the online learning environment differently; while all instructors *Presented* less, this may not have been an intentional practice. For instance, if instructors had to explain to students how to complete their assignments online, then they had to incorporate more *Administering* practices, instead of purposefully intending to reduce their *Presenting* practices. However, as most instructors incorporated more *Guiding* activities, this led to more student participation in their class sessions. Mean percentages for all instructors collapsed COPUS codes can be found in File S11.

### Students who engaged in more W*orking and Talking* behaviors had instructors who implemented more G*uiding* practices in the online learning environment

We found that students were *Working & Talking* more in the online learning environment if their instructor incorporated more *Guiding* practices (Figure 2). For example, in the online learning environment, Marino’s class sessions consisted of mostly *Guiding* practices (50%), and as a result his students *Working & Talking* behaviors were also most of the class time (57%). However, even the instructors who used less *Guiding* practices in the online learning environment had high *Working & Talking* behaviors. For example, during online instruction Patrisia *Guided* less (42%) compared to in-person instruction (50%), but her student *Working & Talking* behaviors remained almost the same in online instruction (51%). Therefore, we interpreted that students had more opportunities in the online learning environment to be able to participate in *Working & Talking* behaviors even if they weren’t facilitated by the instructor, such as using the chat to solve problems amongst multiple students. Mean percentages for all student collapsed COPUS codes can be found in File S12.

### Macie spent more class time on *clicker questions, follow-up,* and *moving and guiding* in the online learning environment

To showcase how online instruction and online functions shifted the use of specific teaching and learning practices, we took a closer look at Macie’s individual COPUS and E- COPUS codes (Figure 3). Looking at Figure 3A, we see that Macie spent more class time on five of the six *Guiding* codes in the online learning environment compared to in-person instruction: *follow-up, posing a question, clicker question, answering questions*, and *moving and guiding*. During in-person instruction, Macie spent about 6% of her class time on c*licker questions,* while during online instruction Macie spent 16% of her class time on c*licker questions.* This could be because clicker questions were already formatted to be administered online; therefore, it was a somewhat easy *Guiding* practice for instructors, such as Macie, to implement in the online learning environment. Following c*licker question*, *follow-up* was the next *Guiding* code that Macie spent more class time on in the online learning environment. During in-person instruction, Macie spent an average of 4% of class time on *follow-up*, while during online instruction she spent an average of 12% of class time on *follow-up.* Since Macie incorporated more *clicker questions* in the online learning environment, she also spent more time explaining student responses and correct answers, which is considered *follow-up*. Penultimately, Macie spent 3% of the class time *moving and guiding* students in active learning activities during in-person instruction, but online she spent an average of 11%. During *clicker questions*, Macie spent more time *guiding* her students in this active learning activity by providing them hints and tips to solve the problem, which is considered a *moving and guiding* behavior. We interpret this to mean that Macie took advantage of the online learning environment and purposefully shifted her teaching practices from teacher-centered to more student-centered approaches. Mean percentages for Macie’s individual instructor COPUS codes can be found in File S13.

### Macie’s students spent more class time engaged in *individual thinking, answering questions,* and *whole class discussions* in the online learning environment

Looking at Figure 3B, Macie’s students also spent more class time on *Working & Talking* behaviors during online instruction. More specifically, Macie’s students spent more time doing *individual thinking,* participating in *whole class discussions,* and *answering questions* online. During in-person instruction, Macie’s students spent an average of 4% of class time on *individual thinking*, while during online instruction students spent an average of 20% class time on *individual thinking*. This increase in *individual thinking* can easily be explained by the increased use of *clicker questions.* Additionally, during in-person instruction, Macie’s students spent an average of only 1% of class time on *whole class discussion,* while during online instruction they spent 14%. These discussions could have naturally emerged as Macie incorporated more *Guiding* behaviors, such as *follow-up,* or as students had more modalities available to engage in *whole class discussions,* such as the chat. Overall, we interpret that students in Macie’s online class sessions had more opportunities to engage in active learning activities as a result of the new online functions, such as polling and the chat. Mean percentages for Macie’s individual student COPUS codes can be found in File S14.

## Discussion

The transition to ERT revealed innovative teaching practices utilizing different online functions to engage students in the online learning environment. Our findings suggest that ERT forced instructors to re-think and re-design their in-person teaching practices to engage students in the new learning environment. We developed and validated E-COPUS to measure these teaching practices in the synchronous online format. More specifically, we adapted six instructor COPUS code descriptions to better represent the observed online teaching practices: *moving and guiding, one-on-one, administering, clicker questions, posing questions,* and *instructor answering questions.* Moreover, we adapted six student COPUS code descriptions to better represent the observed online learning behaviors: *student answering questions, whole class discussion, group clicker question, group worksheet, other group work,* and *student question*. Our instructors did not demonstrate any new teaching or learning behaviors in the online learning environment; instead, they adjusted to their in-person teaching practices by utilizing online functions, such as breakout rooms, polling, the chat, and more. For example, our instructors did not stop *moving and guiding* students once we transitioned to online instruction; but rather, they adapted to the online environment and utilized the breakout rooms function and moved between breakout rooms to mimic *moving and guiding* in the in-person environment. Similarly, Rupnow et al. (2020) found that their instructors adapted their in-person teaching practices to fit the online learning environment, such as drawing using an annotation function on Microsoft Teams instead of drawing on a physical whiteboard in-person.

### Instructors who *Guided* more in the online environment had increased student *Working and Talking* behaviors

Regarding our application of E-COPUS, our data showed that our six instructors adapted their teaching practices in the online learning environment by spending less time on *Presenting* behaviors and spending more time on *Administering* behaviors (File S11). We can infer that instructors had to spend more time going over *Administrative* tasks and classroom logistics with students, such as explaining where to find assignments online or assigning breakout rooms, which explain why there were less *Presenting* codes for all instructors in the online environment. Similarly, Perets et al. (2020) assessed student learning experiences before and after ERT through surveys and found that student engagement decreased when an instructor was lecturing after transitioning to ERT. Based on the results of Perets et al. (2020), we can infer that instructors *Presenting* less in the online learning environment allowed the students we studied to be more engaged in other classroom activities, such as individual and group work in breakout rooms, in chat, and via polling. Additionally, some adapted more *Guiding* practices in the online learning environment; for example, *Guiding* practices were higher for Macie, Paige, Daphne, and Marino after transitioning to synchronous online instruction. While we did not analyze the specific *Guiding* practices implemented by each instructor, we can infer that the online environment gave them opportunities to incorporate more active learning practices, especially with the addition of the online functions. Our findings contradict previous studies concluding that ERT had a negative impact on student engagement (Perets et al., 2020; Reinholz et al., 2020) and loss of interaction and communication between instructors and their students (Petillion & McNeil, 2020).

Regarding student behaviors, we found that the four instructors who *Guiding* more during online instruction engaged their students in more *Working & Talking* behaviors. Similarly, Venton and Pompano (2021b) found that chemistry instructors implementing active learning activities were highly successful in engaging students and maintaining student attendance in the online learning environment. More specifically, they found that when students were assigned to breakout rooms, they were more likely to turn on their cameras and engage with the material and their peers because they knew their peers were depending on them (Venton & Pompano, 2021a). Similar to in-person findings in Kranzfelder, Lo, et al. (2019), our data suggest that if instructors implemented more *Guiding* activities, then students were most likely engaged in more *Working & Talking* behaviors in the online learning environment. Our results are important to show instructors what online practices effectively engage their students in active learning activities and what online functions can be used to support the implementation of these activities.

### Macie incorporated more *Guiding* and students engaged in more *Working and Talking* in the online learning environment

Based on our E-COPUS findings, Macie spent more class time on *Guiding* behaviors, including *clicker questions, follow-up,* and *moving and guiding*. Furthermore, she spent less time *lecturing* in the online learning environment compared to in-person. During her online class sessions, instructor Macie spent more time *moving and guiding* students in an active learning activity by providing hints to students as they completed the activity and spent more time *following-up* with students after *clicker questions* than she did in-person, which led students to spending more time *individually thinking* online. Our findings are consistent with Baldock et al. (2021) who also implemented clicker questions in the online learning environment and found them useful in helping instructors not only provide prompt feedback to students, but also to keep their engagement high during the class session. Therefore, we can conclude that the online clicker questions, or the polling function, were effective tools in engaging students in the online learning environment.

Macie’s students also spent more time engaged in *Working & Talking* behaviors online. Macie’s students frequently participated in *whole-class discussions* and *answered questions*, which could be attributed to the online chat function. Similarly, Tan et al. (2020) found that in their virtual chemistry classroom, the chat was an essential function which allowed their students to engage in discussions at any time during the class session instead of waiting for the instructor to pause and allow students to participate in discussion. Therefore, as instructors begin to transition back to in-person or hybrid instruction, they may consider how implementing online functions which promote student engagement, such as the chat, can be applied in other teaching modalities, such as in-person or hybrid instruction.

Based on our findings, Macie’s data illustrated how the online learning environment and online functions can provide an opportunity for a classroom to include more *Guiding* and *Working & Talking* behaviors. In contrast to previous studies which discovered that instructors used more instructor-centric, lecture-based teaching practices during ERT (Erickson & Wattiaux, 2021), we found that our instructors implemented more student-centric, active learning teaching practices. Although we did not study instructors’ perceptions of teaching and learning in this study, perhaps our instructors’ beliefs, values, changed during ERT. Many of our instructors could have decided that the transition to ERT served as an opportunity to test out new, more student-centered teaching practices, like individual thinking, clicker questions, or small group work. During the transition to and the continuation of ERT, our institution offered educational development opportunities to support instructors as they transitioned to online instruction with the implementation of evidence-based teaching practices, like active learning for large enrollment classes. These efforts could have helped our instructors become more skillful and motivated to implement their teaching practices as observed during ERT.

### Applications of E-COPUS

The innovative teaching practices and online functions instructors adopted to engage their students in the online learning environment will persist as instructors teach in different learning environments. E-COPUS results can benefit instructors, institutional assessment programs, and biology education researchers in many ways. First, if instructors have previously received in-person COPUS data, then they can compare their two sets of data to see if and how their teaching practices have changed between in-person and online instruction. For example, we compared instructor Macie’s in-person and online COPUS data to see that she incorporated more *Guiding* activities in the online learning environment, which led to her students engaging in more *Working & Talking* behaviors. Whether this was planned or not, we were able to see the student behaviors that emerged because of the *Guiding* activities she implemented. When Macie returns to in-person instruction, she may consider how she can continue to use the same *Guiding* practices that she utilized online in the in-person learning environment.

Additionally, instructors can use their E-COPUS data to explore what online functions or active learning practices engaged their students in the most *Working and Talking* behaviors. For example, in Macie’s online learning environment, the chat was crucial to allow students to engage in frequent *whole class discussions* with their peers. By observing this in the E-COPUS data, Macie now knows which teaching practices effectively engaged her students in *Working & Talking* behaviors in the online learning environment and may consider implementing a live chat in her in-person classroom. Instructors across all universities can benefit from E-COPUS data as it provides them with understanding which online teaching practices engaged students in the most *Working and talking* behaviors and will allow them to consider if any effective online practices could be implemented during in-person instruction.

As some universities begin to return to in-person instruction, Hybrid, or Hybrid-Flexible (HyFlex) instruction (Keiper et al., 2020; Kohnke & Moorhouse, 2021; Miller et al., 2020), E- COPUS and COPUS can be used to record both the online and in-person teaching practices. By having a standard protocol for both the in-person and online environment, it will allow for consistent classroom observations between the two class formats. If instructors decide to incorporate online functions into the in-person learning environment, COPUS and E-COPUS can be implemented together to document instructor and student behaviors. We hope this tool will support instructors in understanding and improving their own teaching practices, as well as provide researchers with a tool that can be used consistently in the online environment. Furthermore, if universities continue or revert to online instruction in the future, then instructors can refer to their E-COPUS data to determine what practices were effective for them in the past, as well as where they can improve. Overall, E-COPUS can be applied to 1) understand the teaching and learning behaviors instructors adapted during online instruction, 2) how these behaviors changed between in-person and online instruction, and 3) make decisions on what teaching practices to implement when instructors return to in-person/hybrid instruction, or if instructors return to online instruction in the future.

## Limitations and Future Directions

We acknowledge that there are several factors that limit our study, providing opportunities for future studies. First, we conducted a convenience sample at one MSI, UC Merced; therefore, our results have limited generalizability. Our selected participants were teaching introductory chemistry and biology courses based on the focus of the larger grant-funded studies, so we did not employ a systemic approach to ensure even distribution of faculty and students across STEM disciplines at our institution or other institutions. In the future, it would be interesting to collect E-COPUS data across several universities, especially other MSIs, to determine if the teaching and learning practices at our institution are similar to others.

In addition, we developed E-COPUS while observing instructors using Zoom during synchronous online instruction; therefore, we did not examine if there were differences in teaching and learning practices across different meeting software programs, such as Skype or Google Meet. We recommend future studies utilize E-COPUS to document online behaviors with other software programs to see if new teaching or learning behaviors emerge, and if our current code descriptions are applicable outside of the Zoom meeting software program. Furthermore, we hope that future studies will utilize E-COPUS to document how instructors incorporate newfound online functions, such as the chat, during in-person instruction.

As we continue to assess the online learning environment, E-COPUS could be complemented by pairing it with other tools to study other variables which influence online instruction. Teacher discourse moves, or the conversational strategies used by instructors to encourage student engagement in science content (Kranzfelder et al., 2020; Warfa et al., 2014), has not yet been studied in the online learning environment. Observing instructor discourse alongside instructor behaviors can reveal the quality of active learning strategies used by instructors in the online learning environment. For example, instructors may be using student-centered, *Guiding* teaching practices, but taking a teacher-centered, *authoritative* discourse approach with their students (Kranzfelder et al., 2020). By pairing E-COPUS with discourse protocols, such as the Classroom Discourse Observation Protocol (CDOP) (Kranzfelder, Bankers-Fulbright, et al., 2019), instructors can assess if their teaching practices align with how they are talking to their students.

Finally, we focused on the teaching and learning practices at an MSI, but we did not study how the different student demographics were impacted by changes in the teaching practices because of the transition to ERT. In the future, it would be relevant to examine aspects of equity and inclusion as well as power dynamics in the online environment by taking a closer look at student behaviors. Based on recent studies, Barber et al. (2021) found that first-generation and underrepresented minority students were more likely to have limited access to the internet and computers compared to their white counterparts, suggesting that flexibility on policies and assignments would create a more equitable online learning environment. Also, Lee and Mccabe (2021) found that male students dominated in-person discussions in science courses compared to their female counterparts. Furthermore, they found that male students frequently spoke without raising their hands and used assertive language when speaking (Lee & Mccabe, 2021). The online learning environment is unique in that it allows for students to participate using both the messaging function and verbally, which may lead to female students participating as frequently as their male counterparts. In the future, we recommend documenting who is talking and students’ modes of communication during online, hybrid, and/or in-person instruction.

## Acknowledgments

We would like to thank all the instructors who welcomed us into their classrooms. We would also like to thank all the SATAL interns for their contributions to COPUS data collection and brainstorming E-COPUS code descriptions: Andrea Alvarez Gallardo, Ashley Argueta, Ayooluwa Babalola, Gurpinder Bahia, Avreen Bal, Guadalupe Covarrubias-Oregel, Sandy Dorantes, Dafne Garcia, Jesus Lopez Lopez, Matias Lopez, Stephanie Medina, Andrew Olmeda Juarez, Sara Patino, Monica Ramos, Simrandip Sandhu, Kim Ta, Christian Urbina, Shaira Vargas, Abrian Villalobos, Riley Whitmer, and Isabella Woodruff. Special thanks to members of the Schuchardt and Warfa research teams at the University of Minnesota for their valuable input as our expert panel feedback and manuscript revisions.

