## Supplemental Materials for "Breakout Rooms, Polling, and Chat, Oh My! The Development and Validation of Online COPUS"

Table of Contents

**Supplemental File 1.** E-COPUS Development Flowchart……………………………………...…………2

**Supplemental File 2.** E-COPUS Training Components ………………….…….…………........………....3

**Supplemental File 3.** Instructor COPUS Frequently Asked Questions …….……………………….…….4

**Supplemental File 4.** Student COPUS Frequently Asked Questions …………………….........………….8

**Supplemental File 5.** In-person COPUS Training Inter-Rater Reliability Calculations Among	Coders………………………….…………………………………………………………………13

**Supplemental File 6.** E-COPUS Training Inter-Rater Reliability Calculations Among	Coders…………………..……………………………………………………………………..….14

**Supplemental File 7.** Original E-COPUS Scheme and Expert Panel Feedback…….…………..….……15

**Supplemental File 8.** Matching Activity Responses………...…………………………………………...23

**Supplemental File 9.** E-COPUS Instructor Coding Scheme with Inclusion and Exclusion	Criteria...……………………………………….……………………………………....……...….25

**Supplemental File 10.** E-COPUS Student Coding Scheme with Inclusion and Exclusion

Criteria…………………….…………….………………….…………………………………….27

**Supplemental File 11.** Mean Percentages of Collapsed Instructor COPUS Codes by Instructor and Class	Format..........……………………………………………………………….…………………......29

**Supplemental File 12.** Mean Percentages of Collapsed Student COPUS Codes by Instructor and Class	Format …........................................................................................................................................30

**Supplemental File 13.** Mean Percentage of Two-Minute Time Intervals for Macie’s Individual Instructor	COPUS Codes…………………………….………………………………………........................31

**Supplemental File 14.** Mean Percentage of Two-Minute Time Intervals for Macie’s Individual Student	COPUS Codes ……………………………...………………………….…………........................32

**Supplemental File 1.** E-COPUS Development Flowchart. The development process of E-COPUS consisted of in-person reliability training and observations, online observations with discussion online teaching and learning behaviors to discuss code descriptions, validation, and online reliability training and observations.

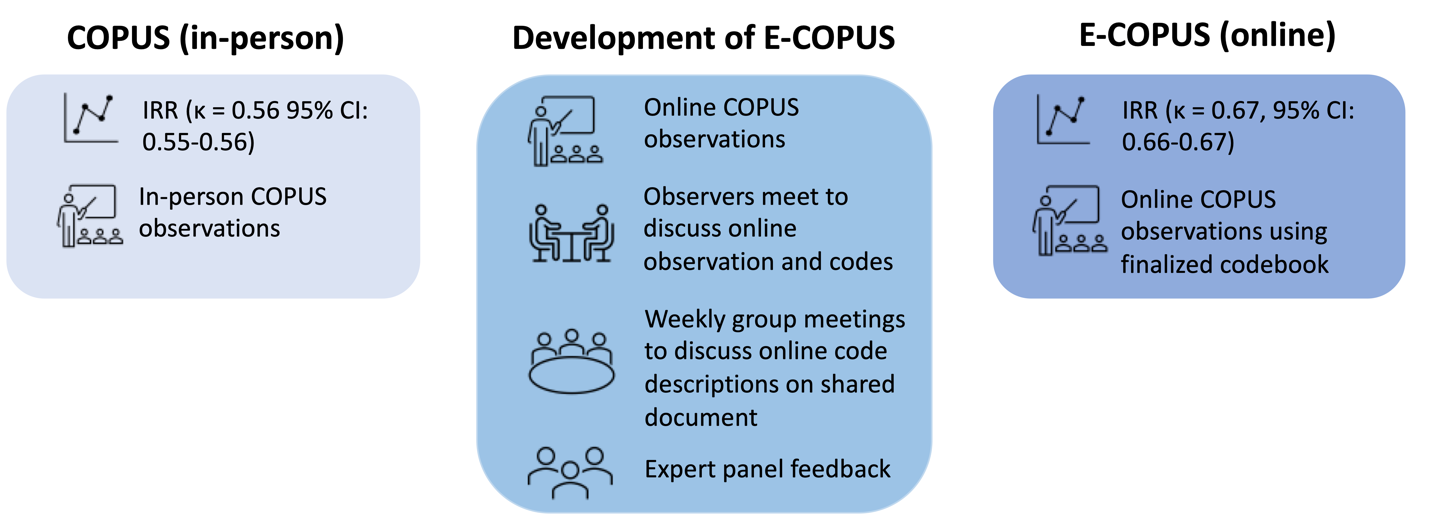

**Supplemental File 2**. E-COPUS Training Components. Training process is used for all new observers and includes individual preparation materials, an interactive activity, pre and post coding activities, an in-depth coding activity following Smith et al 2013 utilizing lecture recordings from our home institution, and assessment to gauge understanding of COPUS code descriptions and protocol.

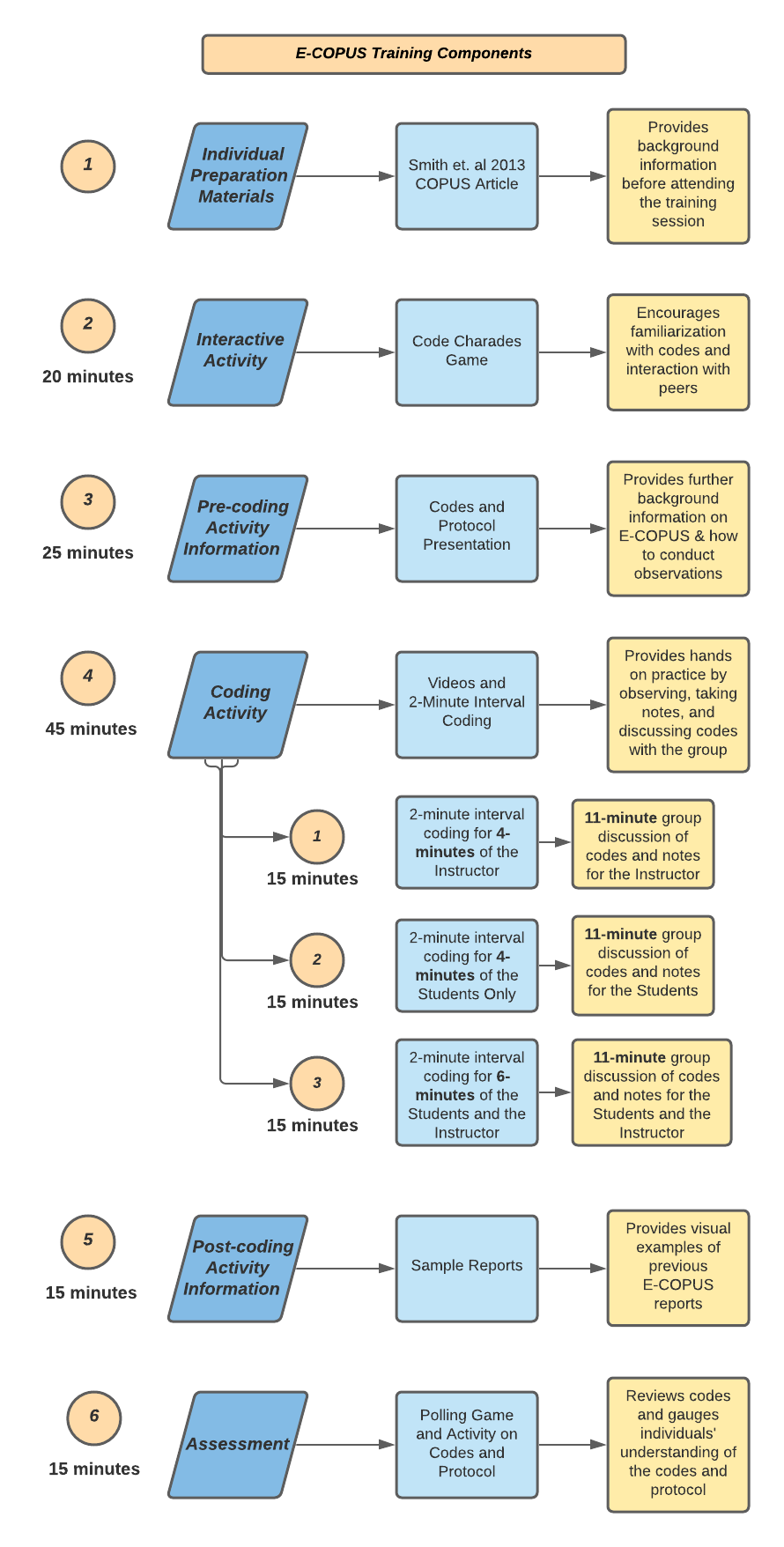

**Supplemental File 3.** Instructor COPUS Frequently Asked Questions. Includes original definitions of each code along with further descriptions, common pairings, and frequently asked questions for different scenarios that commonly occur in the classroom.

| **Code** | **Instructor is Doing** |
| --- | --- |
| **Lec** | **Quick Definition:** Lecturing (presenting content, deriving mathematical results, presenting a problem solution, etc.)  **Description:**   - Includes discussion on learning objectives, exam content, or other course content information. - Transitions from **FUp** to **Lec** may be hard to notice.     **FAQ:**  **Q:** Do we mark **Lec** when the instructor is explaining a clicker question?  **A:** No, if the instructor poses a clicker question and then explains the question, that would be **FUp**. However, if the instructor is going over information related to the problem *before* introducing the clicker question, then it would be **Lec**. |
| **RtW** | **Quick Definition:**  Real-time writing on board, document projector, etc. (often coded simultaneously with Lec)  **Description:**   - Includes the instructor writing or drawing on a device projected to the whole class.     **FAQ:**  **Q:** If the instructor is writing on a piece of paper and does not show it to students until after they are done writing, do we still mark **RtW**?  **A:** Yes, mark **RtW** while the instructor is writing, because they are still writing in real time. However, when the instructor is just showing the piece of paper to students, you do not. |
| **FUp** | **Quick Definition:**  Asking a follow-up question, providing answers/feedback on clicker questions, student responses, or other activity to the entire class.  **Descriptions:**   - **FUp** is coded when the instructor is asking a student a follow-up question on their response to an activity. - **FUp** is coded when the instructor shows results for a clicker question, explains the correct or incorrect answers, or asks for volunteers to share their answers. - If the instructor asks students a follow-up question, you can pair **FUp** with **PQ**.     **FAQ:**  **Q:** When an instructor asks for volunteers to share their answers for a question or activity, is that PQ or FUp?  **A:** This would be both **PQ** and **FUp** since it was related to a previous question or activity.    **Q:** Is it FUp when the instructor says “good job” to the student after they answered a question since it is technically feedback?  **A:** This would not be FUp because it is not providing answers or feedback on the  clicker question or response.    **Q:** Does **FUp** have to be about a clicker question or activity?  **A:** No, **FUp** can be coded when the instructor expands on a student’s response, asks a follow-up question, or potentially any other back and forth conversation between student(s) and instructor. |
| **PQ** | **Quick Definition:** Posing non-clicker questions to students (non-rhetorical) and waiting for students to respond.  **Description:**   - Instructor is asking students a question that requests a response. - If an instructor asks a question but continues to talk (they did not wait for a response), then you would not code **PQ**, it would just be **Lec**.     **FAQ:**  **Q:** If the instructor asks a question about homework or assignments, do we code **PQ** or **Adm**?  **A:** If the instructor is asking students administrative questions about tests, assignments, etc., you can pair **PQ** with **Adm**. |
| **CQ** | **Quick Definition:** Asking a clicker question (mark the entire time the instructor is using a clicker question, not just when first asked) (Can be paired with **Ind** or **CG**)  **Description:**   - Code **CQ** for the whole time it is being administered.     **FAQ:**  **Q:** When an instructor poses a question to the whole class, but provides multiple choice options, would that be **PQ** or **CQ**?  **A:** Since students are not submitting an answer using a clicker and they still must share their answers verbally, it would be **PQ.**    **Q:** If the instructor uses clicker questions as a test, do we code **CQ** or **Adm**?  **A:** Since it is a test, you can pair **Adm** and **CQ** to show that the instructor is administering a test or quiz, but that it’s in clicker question format. |
| **AnQ** | **Quick Definition:** Listening to and answering student questions *with entire class listening.*  **Description:**   - Instructor answers a student’s question in front of the whole class.     **FAQ:**  **Q:** What if an instructor goes to answer a student’s raised hand (**SQ**)?  **A:** You can code that instance as **MG, 1o1,** and **ANQ**. Since the instructor does answer the student’s question, we want to acknowledge that with **AnQ**. However, since it’s not with the entire class listening, we pair it with **MG** and **1o1**. |
| **MG** | **Quick Definition:** Moving through class guiding ongoing student work during active learning task.  **Description:**   - The instructor is moving throughout the classroom observing student work, individual or group. - The instructor does not have to talk to students to code **MG** - **MG** is most paired with **1o1**, when the instructor stops to talk with an individual student or student group.     **FAQ:**  **Q:** Do I code **MG** when the instructor is moving back and forth in front of the class?  **A:** No. What is key about **MG** is that the instructor is moving throughout the whole class, so just walking back and forth would not be considered **MG**. |
| **1o1** | **Quick Definition:** One on one extended discussion with one or a few individuals, giving undivided attention to one or a group of student**s** (can be along with **MG** or **AnQ**).  **Description:**   - The instructor provides individual attention to students. - This can be paired with **AnQ** or **MG** if the instructor is answering a student question or spending individual time with more than one group of students.     **FAQ:**  **Q:** If I can hear what the instructor is saying, for example if they ask individual students a question, should I code it?  **A:** No, try to be as objective as possible. You are trying to capture a holistic view of the classroom, not what the instructor talks about with students individually. Pretend you cannot hear what they are saying and focus on the practices instead of discourse.    **Q:** How long do instructors have to stay with students to code **1o1**?  **A:** If the instructor just passes by and says a sentence to a student, it would not be **1o1**. If an instructor spends a prolonged amount of time with a student discussing something, then you can code **1o1**. We usually consider anything over 30 seconds as **1o1**. |
| **D/V** | **Quick Definition:** Showing or conducting a demo, experiment, simulation, video, or animation.  **Description:**   - **D/V** includes GIFs, physical demonstrations done by the instructor, or even music being played. - If the instructor simply holds up an object, it is not considered to be **D/V.**     **FAQ:**  **Q:** Do I code **Lec** with **D/V**?  **A:** No, if the instructor is not talking, you only code **D/V**. If the instructor is giving information *before* playing the video, you can code **Lec**. If the instructor provides information *after* the video, you can code **FUp**. |
| **Adm** | **Quick Definition:** Administration (Assigning homework, returning tests, class announcements/agenda, etc.)  **Description:**   - Administration counts for administrative work, such as discussing the agenda, assigning homework or tests, discussing office hours, etc. - When students are **TQ**, the instructor should be **Adm.** - **Adm** would also include when the instructor is waiting for students to think in a brainstorm activity, such as a think-pair-share. However, if the instructor is moving throughout the class, you can pair **Adm** and **MG.** |
| **W** | **Quick Definition:** Waiting when there is an opportunity for an instructor to be interacting with or observing/listening to student or group activities and the instructor is not doing so.  **Description:**   - **W** does not include when the instructor is waiting for students to answer a question, that is **Adm.** - **W** is for when there is a *missed opportunity* to interact with students, such as during group work. |
| **O** | **Quick Definition:** Other – explain in comments.  **Description:**   - Only use this code if there is a behavior that does not fit within the other codes. |

**Supplemental File 4.** Student COPUS Frequently Asked Questions. Includes definitions of each code along with further descriptions, common pairings, and frequently asked questions for different scenarios that commonly occur in the classroom.

| **Code** | **Students are Doing** |
| --- | --- |
| **L** | **Quick Definition:** Listening to instructor, taking notes, etc.    **Description:**   - Often coded with Lec, D/V, FUp, Adm. - Can be coded while some students are answering questions with the rest of the class listening. - Can be coded along with SQ when a student(s) is giving a presentation or solving a problem in front of the class.     **FAQ:**  **Q:** Do I code L if the instructor is telling anecdotal stories that only somewhat or barely relate to what is happening in class?  **A:** Yes, L includes listening to anecdotal stories only somewhat related to class |
| **Ind** | **Quick Definition:** Individual thinking/problem solving **(individual clicker question, think-pair-share, private messaging the instructor, or individual test/quiz, etc.).** Only mark when an instructor explicitly asks students to think about a clicker question or another question/problem on their own.    **Description:**   - Can be paired with the instructor codes CQ if it is a clicker question or PQ if it is a think pair share or other prompt and the instructor gives students time to work on it. - Do not code as Ind if the instructor asks the group to raise hands to answer a question.     **FAQ:**  **Q:** Do we mark Ind only when first asked?  **A:** No, mark Ind throughout the entire activity.    **Q:** What if they are asked to make a prediction on their own using a clicker?  **A:** If a clicker question is used to make a prediction on their own, mark it both as Ind and Prd. You only mark CG when they are discussing clicker questions in groups of 2 or more.    **Q**: When a question is presented but the instructor doesn’t allow the students to immediately start working on it, do you code Ind when the question is asked or when they are allowed to work on it?  **A:** Only code Ind once the students are allowed to work on it |
| **CG** | **Quick Definition:**  Discussing clicker question in groups of 2 or more students**.**    **Description:**   - Can be used with Ind if students are asked to think individually before completing a group clicker question. - Code as OG if they are using a non-electronic voting system. - If a clicker question is used to make a prediction in groups of 2 or more students, mark it both as CG and Prd.     **FAQ:**  **Q:** Do all polling activities count as clicker questions?  **A:** Yes, mark CG if students are working on a poll question with a group of students.    **Q:** When an instructor asks a question and doesn’t allow the students to immediately start working on it, do you code CG when the question is asked or when they are allowed to work on it?  **A:** Only code CG once the students are allowed to work on it |
| **WG** | **Quick Definition:** Working in groups of 2 or more students on worksheet activity.  **Descriptions:**   - Only use this code for actual worksheet activities that were handed out during class. This includes worksheets that were handed out in previous classes but were not yet finished. - Instructor codes would be PQ since it is not a clicker question and Adm since the instructor is waiting for students to complete the activity.       **FAQ:**  **Q:** Do you still code WG if the class does not turn in the worksheet at the end of class?  **A:** Yes, you still code WG even if the worksheet isn’t turned in at the end of class.    **Q:** What are the distinguishing factors between WG and OG?  **A:** WG should only be coded if a sheet with questions is handed out for the student to work on, otherwise the code should be OG. For example, an instructor having students working problems from a book and writing down the answers on their own paper would count as OG since a worksheet wasn’t handed out. |
| **OG** | **Quick Definition:** Working in groups of 2 or more students on other assigned group activity, such as responding to instructor question.    **Description:**   - This could include various activities such as writing in course notebooks, answering questions on online modules, answering a group prompt or question, discussing an activity with a group etc. Try to list the activity in the notes section. - If the activity involves making a prediction, code Prd at the same time. - Code MG if the instructor moves from group to group.     **FAQ:**  **Q:** Are questions that are assigned in groups considered OG?  **A:** Yes, but only if it is not a group clicker question. If it is a clicker question, code CG. |
| **AnQ** | **Quick Definition:** Student answering a question posed by the instructor with the rest of the class listening.    **Description:**   - Code AnQ for instructor side when TA answers a student question. - Can be paired with WC if more than one student answers a question. - Can be paired with Ind if students think about the question before answering. - Use this code for both verbal answers and when students are asked to raise their hands.     **FAQ:**  **Q:** If the instructor asks students to raise their hands when they are done with an activity is this AnQ?  **A:** Yes, non-verbal responses such as raising their hand or nodding their head to an instructor’s question or comment is also AnQ. |
| **SQ** | **Quick Definition:** Student asks a question.    **Description:**   - Do not code when students’ answer to a question seems unsure so it sounds like a question. This would still be AnQ. - This is used when students ask questions in front of the large group. - A student could be asking a question to the instructor or to another student.     **FAQ:**  **Q:** Do raised hands during group work count as a SQ?  **A:** Yes, you count a raised hand as SQ. |
| **WC** | **Quick Definition:** Instructor poses a question or facilitates a whole class discussion in which 2 or more students answer the rest of the class is listening.    **Description:**   - Can be paired with AnQ and SQ. - Coded when multiple students are answering or providing their thoughts on one question. - The instructor may have an active (e.g., providing new questions/responses) or passive (e.g., selecting speakers) role in the discussion.     **FAQ:**  **Q:** Does the instructor have to explicitly say that the whole class can discuss for it to be a whole class discussion?  **A:** No, if the instructor asks a question or poses a prompt and all students can participate even if not explicitly stated, WC can be coded. |
| **Prd** | **Quick Definition:** Predicting the outcome of a demo/experiment.    **Description:**   - Only code when the instructor explicitly tells students to “predict” - This could include time thinking on their own or to raise their hand based on what they think will occur (in this special case also code Ind) but only if the instructor tells them to “predict.” - Can be paired with D/V (demo, experiment, or video) - Can be paired with Ind, CG, OG, or WG if students are predicting and thinking about a question.     **FAQ:**  **Q:** When an instructor asks students to make a prediction but doesn’t allow the students to immediately start working on it, do you code Prd when the instructor asks or when they are allowed to work on it?  **A:** Only code Prd once the students are allowed to work on it.    **Q:** If the instructor asks the students to make a new prediction or refine the one, they already made while an experiment or demonstration is happening, do I code Prd and D/V at the same time?  **A:** Yes, if the instructor has already begun the experiment and asks the students to make new predictions than they should be coded simultaneously. |
| **SP** | **Quick Definition:** Students facilitate a presentation, solve a problem, or explain a figure while the rest of the class is listening.    **Description:**   - Code when students have some time to think about something before presenting it to the class even if the time is brief. - This could include a pre-assigned student presentation or an occasion where the instructor asks a student to “take over” the class to teach a concept or to present a solution to a problem (or other activity) to the entire class. - Code with L for the rest of the class   **FAQ:**  **Q:** Do we code SP when a student volunteer is asked to annotate on a slide in response to a previously posed question or activity?  **A:** Yes.    **Q:** What do we code if multiple students are asked to go over a problem in front of the entire class after they were asked to prepare their answers?  **A:** SP, AnQ, WC. |
| **TQ** | **Quick Definition:** Test or quiz    **Description:**   - Pair with Ind, CG, or OG to describe what type of quiz it is. - Code if the question is described as a test or quiz question.     **FAQ:**  **Q:** Do I only code TQ when the test or quiz is first assigned?  **A:** No, code TQ throughout the entire activity. |
| **W** | **Quick Definition:** Instructor late, working on fixing technical problems, instructor otherwise occupied, etc.)    **Description:**   - Do not mark quick transitions. - Mark when the instructor explicitly says they will wait for students to answer a question.     **FAQ:**  **Q:** If the instructor assigns a test or quiz and is waiting for the student to answer is this W?  **A:** No, since the instructor is administering the test or quiz, you would code Adm. |
| **O** | **Quick Definition:** Other- explain in comments.    **Description:**   - Can be used if the instructor is interrupted during class. - Only use when another code does not fit the situation. If this code is used many times revise your codes to see if the scenarios can fit in another code. |

**Supplemental File 5.** In-person COPUS training - inter-rater reliability among coders. Data was taken collected from a 30-minute class session that was coded by 16 coders.

| **Instructor** | **Class** | **Coder pairs** | **No. of minutes** | **Fleiss’ Kappa** | **Confidence Intervals** | |
| --- | --- | --- | --- | --- | --- | --- |
| **Code** | **Session** |  |  |  | **Lower** | **Upper** |
| **144** | 4-Nov-19 | All 15 coders | 30 | 0.55 | 0.55 | 0.56 |

| **Supplemental File 6.** E-COPUS training - inter-rater reliability calculations among coders. Data was collected from a 30-minute class session that was coded by 23 coders. | | | | | | |
| --- | --- | --- | --- | --- | --- | --- |
| **Instructor** | **Class** | **Coder pairs** | **No. of minutes** | **Fleiss’ Kappa** | **Confidence Intervals** | |
| **Code** | **Session** |  |  |  | **Lower** | **Upper** |
| **147** | 6-Jan-21 | All 23 coders | 30 | 0.67 | 0.66 | 0.67 |

**Supplemental File 7**. Original E-COPUS Scheme and Expert Panel Feedback. Original COPUS Codes and descriptions along with Online COPUS code descriptions and notes that were collected during the expert panel feedback.

| **Instructor Doing** | **COPUS Code** | **Original COPUS Code Description** | **In-person COPUS Code Description Clarifications** | **Online COPUS Code Description** | **Expert Panel Feedback (Questions, comments, concerns, etc.)** |
| --- | --- | --- | --- | --- | --- |
|  | Lecturing (Lec) | Lecturing (presenting content, deriving mathematical results, present a problem solution, etc.) | Lecturing (presenting content, deriving mathematical results, presenting a problem solution, **going over learning outcomes,** etc.) | No change | Doesn’t seem to be a new change, but it is common that it can be interpreted in admin. In publication, explain the reasoning for the change. (Learning outcomes assumed to be under admin) |
|  | Real-time Writing (RtW) | Realtime writing on board, doc. projector, etc. (often checked off with Lec) | No change | Real-time writing on board, doc, projector, **annotating lecture slides, or writing notes on a piece of paper and showing them to the class** (often checked off with Lec)**.** | Some professors have a whiteboard and show the whiteboard on their camera, Zoom has an online whiteboard, some professors set up rigs with papers, google docs, some faculty annotate lecture slides are in person, **chat?** Putting a link in the chat from the slide? Provide rationale |
|  | Demo or Video (D/V) | Showing or conducting a demo, experiment, simulation, video, or animation | No change | No change | N/a |
|  | Follow-up (FUp) | Follow-up/feedback on clicker question or activity to entire class | **Asking a follow-up question,** providing answers/feedback on clicker questions, **student responses,** or other activity to entire class. | No change | Probing could be after a response, “explain your reasoning” or expand on what they said, explain differences between follow-up and probing. It’s about the timing- after the activity, follow-up. |
|  | Posing a question (PQ) | Posing non-clicker question to students (nonrhetorical) | Posing non-clicker question to students (non-rhetorical) **and waiting for students to respond.** | Posing non-clicker question to students (non-rhetorical) **using the microphone or using the chat function and waiting for students to respond.** | Change chat to messaging  Microphone- verbal? |
|  | Clicker question (CQ) | Asking a clicker question (mark the entire time the instructor is using a clicker question, not just when first asked | Asking **and administering** a clicker question (mark the entire time the instructor is using a clicker question, not just when first asked) (Can be paired with Ind or CG) | Asking **and administering** a clicker question **or online poll** (mark the entire time the instructor is using a clicker question, not just when first asked) (Can be paired with Ind or CG) | Paired with ind or CG. |
|  | Answering questions (AnQ) | Listening to and answering student questions with the entire class listening | No change | Listening to and answering student questions **using the microphone or using the chat function,** with the entire class listening | N/a |
|  | Moving and guiding (MG) | Moving through class guiding ongoing student work during active learning tasks | No change | **Moving throughout breakout rooms guiding students or guiding students while they are working on a problem or clicker question (hints/working through a problem) using the microphone or chat function** during active learning tasks | Working through a problem is a little less obvious than moving through breakout rooms, doesn’t exactly translate. Module system moving and guiding- a lot of professors and TAs set up modules for the week, and that sets up preparation in the week. Guiding using **probing** **questions.** Differentiate posing, follow-up, and probing questions. |
|  | One on one (1o1) | One on one extended discussion with one or a few individuals, not paying attention to the rest of the class (can be along with MG or AnQ) | One on one extended discussion with one or a few individuals, **giving undivided attention to one or a group of students** (can be along with MG or AnQ) | One on one extended discussion with one or a few individuals, **giving undivided attention to one or a group of students** **in a breakout room or private messaging between instructor and student** (can be along with MG or AnQ) | It can be in chat, what if they @ the student? It’s not in a breakout room. What is the goal of the paper? Fix COPUS to make it more understandable or to adjust to the Online environment? Look at other modifications. Cite that as a reason for making the change, along with the modifications with Online. Why COPUS and not a new codebook? Compare in-person to Online? Should there be two versions of COPUS? What’s the goal? Zoom makes this more likely because of the breakout rooms. **What to do about chat and 1o1. Talking to people with the rest of the class not aware.** |
|  | Administration (Adm) | Administration (assign homework, return tests, etc.) | Administration (Assign homework, return tests, **class announcements, agenda, assign to breakout rooms, etc.),** **when the instructor is waiting for students to answer a non-clicker question (i.e., think-pair-share), or administering a test or quiz** | No change | N/a |
|  | Waiting (W) | Waiting when there is an opportunity for an instructor to be interacting with or observing/listening to student or group activities and the instructor is not doing so | No change | No change | N/a |
|  | Other (O) | Other | No change | No change | N/a |
|  | Listening (L) | Listening to instructor, taking notes, etc. | No change | No change | N/a |
| **Students Doing** | Individual (Ind) | Individual thinking/problem solving. Only mark when an instructor explicitly asks students to think about a clicker question or another question/problem on their own. | Individual thinking/problem solving **(individual clicker question, think-pair-share, private messaging the instructor, or individual test/quiz, etc.).** Only mark when an instructor explicitly asks students to think about a clicker question or another question/problem on their own. | No change | N/a |
|  | Group Clicker Question (CG) | Discuss clicker question in groups of 2 or more students | Discussing clicker question in groups of 2 or more students | Discussing clicker question in groups of 2 or more students **in breakout rooms** | Not everyone uses Zoom! Add online poll? Can’t rely on what is physically possible. But have to use the data we have? Look up other online platforms. Google meet, skype twitch, etc. Whatever your platform is, it should just be a proof of concept. What are the behaviors that are likely to happen when you are teaching remotely? Don’t make it so specific to Zoom. Take the protocol so it doesn’t have to be attached to a specific platform. Many ways to do clicker questions outside of Zoom. (Kahoot, Tophet, etc.) |
|  | Group Worksheet (WG) | Working in groups on worksheet activity | Working in groups **of 2 or more students** on worksheet activity | Working in groups **of 2 or more students** on worksheet activity **in breakout rooms** | N/a |
|  | Other Group Work (OG) | Other assigned group activity, such as responding to instructor question | **Working in groups of 2 or more students on** other assigned group activity, such as responding to instructor question | **Working in groups of 2 or more students on** other assigned group activity, such as responding to instructor question, **in breakout rooms** | N/a |
|  | Answering questions (AnQ) | Student answering a question posed by the instructor with rest of class listening | No change | Student answering a question posed by the instructor **using the microphone function** with the rest of the class listening **or student answering a question posed by the instructor using a reaction function, the annotating function, or the chat function *and* the instructor acknowledges the answer with the rest of the class listening (can be paired with WC or Ind) or student answering other students’ questions using the chat function.** | Reactions are like a group consensus, difference between that and a single answer? More like a poll? If an instructor In-person asked students for a reaction with heads down. Would that still be coded as AnQ? |
|  | Student Question (SQ) | Student asks question | No change | Student asks question **using the microphone or chat function.** | N/a |
|  | Whole class discussion (WC) | Engaged in whole class discussion by offering explanations, opinion, judgment, etc. to whole class, often facilitated by instructor | No change | **Instructor poses a question or facilitate a whole class discussion in which 2 or more students answer using the microphone or chat function while the rest of the class is listening (can be paired with AnQ and SQ) *or* 2 or more students use the annotating function.** | Annotating but not necessarily a discussion about it. If it’s just 2 or more, you’re missing distinguishing features. What about when students talk to each other? Maybe explain that we modified AnQ where students talk to each other also so they can be talking to each other in the whole class discussion. One student vs whole class. **Kelly**: That last code might be another one to mark where Online is going to look different than in person because when you’re observing a whole class in person you can’t capture every time a student answers another students’ question where you can capture more of that using the chat function (although not all because of direct messages.) |
|  | Prediction (Prd) | Making a prediction about the outcome of demo or experiment | No change | No change | N/a |
|  | Student Presentation (SP) | Presentation by student(s) | **No change** | **Students facilitate a presentation, solve a problem, or explain a figure while the rest of the class is listening** | N/a |
|  | Test or Quiz (TQ) | Test or quiz | No change | Test or quiz **(pair with Ind or OG)** | N/a |
|  | Waiting (W) | Waiting (instructor late, working on fixing AV problems, instructor otherwise occupied, etc.) | No change | No change | N/a |
|  | Other (O) | Other – explain in comments | No change | No change | N/a |

**Supplemental File 8**. Matching Activity Responses. The first three scenarios are for instructor code descriptions and the next three scenarios are for student code descriptions were presented to the expert panel. Panelists were asked to match E-COPUS codes to each scenario from a provided list of codes. Each scenario had multiple E-COPUS codes as the correct answers, which are listed under “Correct Answer.” The “Results” were the responses collected from the expert panel. 8 individuals participated in the instructor scenario matching activity, while 6 individuals participated in the student matching activity.

| **The instructor goes to one breakout room and stays guiding a group of students on their assigned clicker question.** | |
| --- | --- |
| **Correct Answer** | **Results** |
| CQ  MG  1o1 | CQ- 8/8  MG- 7/8  1o1- 4/8 |

| **The instructor talks about phenotypes and genotypes. The instructor then asks the class "Does somebody want to raise their hand and help me work through the next part of this?" A student answers and the instructor writes what the student is saying. The instructor stops the student and re-explains to the class what the student is saying. The instructor then asks the student to continue answering. Once the student has finished, the instructor asks the student "How do you get the allele?"** | |
| --- | --- |
| **Correct Answer** | **Results** |
| Lec  Rtw  FUp  PQ | Lec- 8/8  RtW- 6/8  FUp- 6/8  PQ- 8/8  AnQ- 1  1o1- 1 |

| **The instructor waits for students to write down their answers. A student asks in the chat if there is supposed to be a poll. The instructor answers, "just write it down for now".** | |
| --- | --- |
| **Correct Answer** | **Results** |
| AnQ  Adm | AnQ- 6/8  Adm- 8/8  RtW- 2/8  1o1- 2/8 |

| **Students take a moment to think of their response to the question "Why is taxonomy useful?". The instructor calls on a student to answer the question. After the student answers, the instructor asks a student a question about their response. The student answers the instructor's question. The instructor then calls on another student to hear a different answer.** | |
| --- | --- |
| **Correct Answer** | **Results** |
| Ind  AnQ  WC | Ind- 6/6  AnQ 5/6  WC- 6/6 |

| **A student solves a problem in front of the whole class which was assigned to before the class period. The instructor asks him, "Why is the first option not right?". The student answers the question. The student then asks the instructor, "Is this not a C terminal?". The student then continues to explain the question to the class.** | |
| --- | --- |
| **Correct Answer** | **Results** |
| AnQ  SQ  SP | AnQ- 6/6  SQ- 6/6  SP 5/6 |

| **Students go into breakout rooms and are given 10 minutes to work on their group quiz. Students in room 3 share their screen and try to go through the question together.** | |
| --- | --- |
| **Correct Answer** | **Results** |
| OG  TQ | OG 6/6  TQ 5/6 |

**Supplemental File 9**. E-COPUS Instructor Coding Scheme with Inclusion and Exclusion Criteria. Includes E-COPUS codes with descriptions, inclusion and exclusion criteria, and example descriptions of how the codes have applied in the classroom.

| Instructor  Doing | Individual COPUS Code | E-COPUS Code Description | Inclusion Criteria | Exclusion Criteria | Example Description |
| --- | --- | --- | --- | --- | --- |
|  | Moving and guiding (MG) | Moving through **breakout rooms** guiding ongoing student work during active learning task **or guiding students while they are working on a problem or clicker question (hints/working through a problem) using the microphone or messaging function** | The instructor is moving through breakout rooms guiding students through an active learning task by providing hints or steps using the microphone or the messaging function guiding the whole class through an active learning task by providing hints or steps using the microphone or the chat function while students work independently. | The instructor remains in the main room not interacting with students, or guides students through a problem or activity after they have completed it. | *“The instructor gives students hints as to how to approach the problem and underlines the key words of the question.”* |
|  | One-on-one (1o1) | One on one extended discussion with one or a few individuals, giving undivided attention to one or a group of students **in a breakout room** | The instructor remains in a breakout room for more than 30 seconds. | The instructor briefly joins a breakout room but then continues to check in with other rooms, talks with multiple students using the messaging function, or has a back-and-forth conversation with a student in the main room. | *“The instructor stays in one breakout room for the whole activity guiding students.”* |
|  | Posing a question (PQ) | Posing non-clicker question to students (non-rhetorical) **using the microphone or messaging function and** waiting for students to respond | The instructor asks students a question about the course content, poses a question as an activity, or asks students about themselves. | The instructor asks a question and does not wait for students to respond. | *“The instructor asks, ‘What is this one? Type it in the chat’.”* |
|  | Answering questions (AnQ) | Listening to and answering student questions **using the microphone or messaging function** with the entire class listening | The instructor answers a students’ question on the mic or in the chat. | The instructor answers a students’ question during one-on-one time, not in front of the entire class or through private messaging. | *“Instructor answers saying that she is not very sure, she does not know the exact specifics, but it is in the textbook.”* |
|  | Clicker question (CQ) | Asking a clicker question **or online poll** (mark the entire time the instructor is using a clicker question, not just when first asked) | The instructor asks and administrates a clicker question. | The instructor poses a question and provides students the multiple choices, asks students to privately submit answers using the messaging function, or reviews answers to a clicker question. | *“The instructor introduces the clicker question and opens it to the class.’"* |
|  | Administration (Adm) | Assigning homework, returning tests, **class announcements/agenda, assign to breakout rooms, etc.), when the instructor is waiting for students to answer a non-clicker question (i.e., think-pair-share),** **or administering a test or quiz** | The instructor is going through the class agenda, announcements, administering a test or quiz, creating breakout rooms, or staying in the main room while students are in the breakout rooms. | The instructor is administering a clicker question, answering student questions about announcements, or dealing with technical issues. | *“The instructor tells the class that there's no lab or discussion because of the holiday. The instructor tells students they will have a quiz this week.”* |

**Supplemental File 10**. E-COPUS Student Coding Scheme with Inclusion and Exclusion Criteria. Includes E-COPUS codes with descriptions, inclusion and exclusion criteria, and example descriptions of how the codes have applied in the classroom.

| Student Doing | Individual COPUS Code | Online COPUS Code Description | Inclusion Criteria | Exclusion Criteria | Example Description |
| --- | --- | --- | --- | --- | --- |
|  | Answering questions (AnQ) | Student answering a question posed by the instructor **using the microphone function, reaction function, annotating function, or messaging function *and* the instructor acknowledges the answer with the rest of the class listening or student answering other students’ questions using the messaging function** | One or more students answer a question posed by the instructor or answer another students’ question using the microphone function, annotation function, or messaging function in front of the entire class. | The TA answers a student’s question using the microphone or messaging function or students answer the instructor’s question during one-on-one time or through private message. | *“A student is called on and answers the question with his mic. Students listen to the students’ answer.”* |
|  | Whole class discussion (WC) | Instructor poses a question or facilitate a whole class discussion in which 2 or more students answer **verbally, using messaging function, or drawing function while the rest of the class is listening** | Multiple students answer a single question posed by the instructor using the microphone function, messaging or through the annotation function; students build off each other’s responses to a question; multiple students ask questions using the messaging function. | One student answers a singular question from the instructor using the microphone, messaging, or annotation function. | *“15 students share their answers in the chat. Several students also use their microphones and answer: ‘atmosphere’, ‘in lakes’, ‘rivers’, ‘the ocean’, and ‘ice glaciers’ .”* |
|  | Group Clicker Question (CG) | Discussing clicker question in groups of 2 or more students **in breakout rooms or messaging function** | Students discuss or work on clicker questions in a breakout room or using the messaging function. | Students independently work on clicker question. | *“Students go into breakout rooms to discuss their responses to the clicker questions. In breakout room 2, students’ cameras are on, and they discuss the question with each other. In breakout room 3, students do not have their cameras on, but they are sharing the screen with the review questions.”* |
|  | Group Worksheet (WG) | Working in groups of 2 or more students on worksheet activity **in breakout rooms or messaging function** | Students discuss or work on worksheet questions in a breakout room or using the messaging function. | Students independently work on a worksheet. | *“Students have discussion in breakout rooms on the worksheet problems”* |
|  | Other Group Work (OG) | Working in groups of 2 or more students on other assigned group activity, such as responding to instructor question **in breakout rooms in breakout rooms or messaging function** | Students discuss or work on something other than clicker questions in a breakout room or using the messaging function. | Students independently work on an activity. | *“Students go into breakout rooms to have a group discussion. One observer's group does not talk at all.”* |
|  | Student Question (SQ) | Student asks question **using the microphone or messaging function** | Students ask the instructor a question using the microphone or chat function with the rest of the class listening or raises their hand using the raise hand function to ask the instructor a question. | Students ask the instructor a question during one-on-one time or through private message and it is not explicitly stated to the whole class by the instructor. | *“A student asks in the chat, "Do we have discussion today?" and another student points up using "^^" to emphasize the question.”* |

**Supplemental File 11**. Mean Percentages of Collapsed Instructor COPUS Codes by Instructor and Class Format. Collapsed codes include Presenting, Guiding, Administering, and Other (Smith et al., 2013).

| Instructor and Class Format | Presenting | Guiding | Administering | Other |
| --- | --- | --- | --- | --- |
| Jaqueline, In-person | 39.3 | 55.1 | 4.7 | 1.0 |
| Jaqueline, Online | 37.0 | 38.4 | 23.0 | 1.4 |
| Patrisia, In-person | 32.7 | 50.0 | 13.3 | 4.0 |
| Patrisia, Online | 26.7 | 42.2 | 23.0 | 8.1 |
| Macie, In-person | 64.8 | 31.7 | 2.2 | 1.3 |
| Macie, Online | 23.0 | 71.0 | 5.4 | 1.3 |
| Paige, In-person | 68.4 | 24.7 | 5.7 | 1.3 |
| Paige, Online | 51.3 | 37.6 | 8.4 | 2.7 |
| Daphne, In-person | 84.0 | 11.0 | 3.4 | 1.7 |
| Daphne, Online | 54.0 | 34.7 | 10.7 | 1.0 |
| Marino, In-person | 67.8 | 30.9 | 0.7 | 0.7 |
| Marino, Online | 45.0 | 50.3 | 4.3 | 0.5 |

**Supplemental File 12.** Mean Percentages of Collapsed Student COPUS Codes by Instructor and Class Format. Collapsed codes include Receiving, Working & Talking, Assessment, and Other (Kranzfelder, Lo, et al., 2019).

| Instructor and Class Format | Receiving | Working & Talking | Assessment | Other |
| --- | --- | --- | --- | --- |
| Jaqueline, In-person | 41.5 | 55.9 | 0 | 2.5 |
| Jaqueline, Online | 47.6 | 48.4 | 2.4 | 1.6 |
| Patrisia, In-person | 43.6 | 51.9 | 0.0 | 4.5 |
| Patrisia, Online | 44.7 | 50.8 | 3.8 | 1.0 |
| Macie, In-person | 54.3 | 45.7 | 0 | 0 |
| Macie, Online | 35.1 | 62.2 | 0 | 2.7 |
| Paige, In-person | 63.9 | 26.0 | 4.6 | 5.6 |
| Paige, Online | 67.9 | 32.1 | 0 | 0 |
| Daphne, In-person | 78.6 | 14.3 | 0 | 7.1 |
| Daphne, Online | 70.5 | 28.6 | 0 | 1.0 |
| Marino, In-person | 59.0 | 32.0 | 0 | 9.0 |
| Marino, Online | 43.5 | 56.5 | 0 | 0 |

**Supplemental File 13.** Mean Percentage of Two-Minute Time Intervals for Macie’s Individual Instructor COPUS Codes.

| Macie’s Instructor Codes | | |
| --- | --- | --- |
| **Instructor COPUS Codes** | **In-person** | **Online** |
| Lecturing | 0.38 | 0.01 |
| Real-time Writing | 0.25 | 0.26 |
| Demo/Video | 0.02 | 0 |
| Follow-up | 0.04 | 0.12 |
| Posing a Question | 0.11 | 0.19 |
| Clicker Question | 0.06 | 0.16 |
| Answering Question | 0.09 | 0.09 |
| Moving and Guiding | 0.03 | 0.11 |
| One-on-One | 0 | 0 |
| Administration | 0.02 | 0.05 |
| Waiting | 0 | 0 |
| Other | 0.01 | 0.01 |

**Supplemental File 14.** Mean Percentage of Two-Minute Time Intervals for Macie’s Individual Student COPUS Codes.

| Macie’s Student Codes | | |
| --- | --- | --- |
| **Student COPUS Codes** | **In-person** | **Online** |
| Listening | 0.54 | 0.37 |
| Individual | 0.04 | 0.20 |
| Student Question | 0.15 | 0.12 |
| Answering Question | 0.16 | 0.17 |
| Group Clicker Question | 0.06 | 0 |
| Group Worksheet | 0.04 | 0 |
| Whole Class Discussion | 0.01 | 0.14 |
| Prediction | 0 | 0 |
| Student Presentation | 0 | 0 |
| Test or Quiz | 0 | 0 |
| Waiting | 0 | 0 |
| Other | 0 | 0 |
